# Complete mapping of mutations to the SARS-CoV-2 spike receptor-binding domain that escape antibody recognition

**DOI:** 10.1101/2020.09.10.292078

**Authors:** Allison J. Greaney, Tyler N. Starr, Pavlo Gilchuk, Seth J. Zost, Elad Binshtein, Andrea N. Loes, Sarah K. Hilton, John Huddleston, Rachel Eguia, Katharine H.D. Crawford, Adam S. Dingens, Rachel S. Nargi, Rachel E. Sutton, Naveenchandra Suryadevara, Paul W. Rothlauf, Zhuoming Liu, Sean P.J. Whelan, Robert H. Carnahan, James E. Crowe, Jesse D. Bloom

## Abstract

Antibodies targeting the SARS-CoV-2 spike receptor-binding domain (RBD) are being developed as therapeutics and make a major contribution to the neutralizing antibody response elicited by infection. Here, we describe a deep mutational scanning method to map how all amino-acid mutations in the RBD affect antibody binding, and apply this method to 10 human monoclonal antibodies. The escape mutations cluster on several surfaces of the RBD that broadly correspond to structurally defined antibody epitopes. However, even antibodies targeting the same RBD surface often have distinct escape mutations. The complete escape maps predict which mutations are selected during viral growth in the presence of single antibodies, and enable us to design escape-resistant antibody cocktails–including cocktails of antibodies that compete for binding to the same surface of the RBD but have different escape mutations. Therefore, complete escape-mutation maps enable rational design of antibody therapeutics and assessment of the antigenic consequences of viral evolution.

## Introduction

The COVID-19 pandemic has generated urgent interest in antibody therapeutics and vaccines that induce antibodies to SARS-CoV-2. Many of the most potently neutralizing anti-SARS-CoV-2 antibodies target the receptor-binding domain (RBD) of the viral spike protein, often competing with its binding to the ACE2 receptor (Brouwer et al., 2020; Cao et al., 2020; Ju et al., 2020; Liu et al., 2020; Rogers et al., 2020; Seydoux et al., 2020; Wec et al., 2020; Wu et al., 2020; Zost et al., 2020a, 2020b). In addition, anti-RBD antibodies often dominate the neutralizing activity of the polyclonal antibody response elicited by natural infection (Barnes et al., 2020a; Steffen et al., 2020; Weisblum et al., 2020). Both passively-administered and vaccine-induced anti-RBD neutralizing antibodies protect against SARS-CoV-2 in animals (Alsoussi et al., 2020; Cao et al., 2020; Hassan et al., 2020; Rogers et al., 2020; Walls et al., 2020a; Wu et al., 2020; Zost et al., 2020a), and preliminary evidence suggests the presence of neutralizing antibodies also correlates with protection in humans (Addetia et al., 2020).

Determining which viral mutations escape from antibodies is crucial for designing therapeutics and vaccines and assessing the antigenic implications of viral evolution. Escape mutants can be selected by passaging virus expressing the SARS-CoV-2 spike protein in the presence of anti-RBD antibodies in the laboratory (Baum et al., 2020a; Weisblum et al., 2020), and some RBD mutations that alter antibody binding are already present at very low levels in SARS-CoV-2 circulating in the human population (Li et al., 2020). It seems plausible that such mutations could become prevalent over longer evolutionary time, given that the seasonal coronavirus 229E has accumulated genetic variation in its RBD in the last few decades that is sufficient to ablate antibody binding (Wong et al., 2017).

However, current methods to identify SARS-CoV-2 escape mutations by passaging virus in the presence of antibodies are incomplete in the sense that they only find one or a few of the possible escape mutations. Structural biology can more comprehensively define how an antibody physically contacts the virus, but structures are time consuming to determine and still do not directly report which viral mutations escape from antibody binding (Dall’Acqua et al., 1998; Dingens et al., 2019; Jin et al., 1992).

Here we overcome these limitations by developing a high-throughput approach to completely map mutations in the SARS-CoV-2 RBD that escape antibody binding, and apply this approach to 10 human antibodies. The resulting escape maps reveal the extent to which different antibodies are escaped by mutations at overlapping or orthogonal sites, and show that antibodies targeting structurally similar regions sometimes have escape mutations at entirely distinct residues. Furthermore, we show that the escape maps predict which mutations are selected when spike-expressing virus is passaged in the presence of neutralizing antibodies, and can inform the design of antibody cocktails that resist escape. Therefore, complete escape-mutation maps can be used to assess the antigenic consequences of viral genetic variation and the potential for viral escape from specific antibodies or antibody cocktails.

## Results

### A yeast-display system to completely map SARS-CoV-2 RBD antibody-escape mutations

To map antibody-escape mutations in a high-throughput manner, we leveraged a system for expressing conformationally-intact RBD on the surface of yeast cells (Figure 1A). As described previously (Starr et al., 2020), we created duplicate mutant libraries of the RBD from the Wuhan-Hu-1 strain of SARS-CoV-2 that together contained nearly all possible amino-acid mutations in the 201-residue RBD (they contain 3,804 of the 3,819 possible mutations, with >95% present as single mutants). Each yeast cell carries a short 16-nucleotide barcode that identifies the RBD mutant it expresses, enabling us to rapidly characterize the composition of the RBD mutant libraries via deep sequencing of the DNA barcodes.

**Figure 1.**
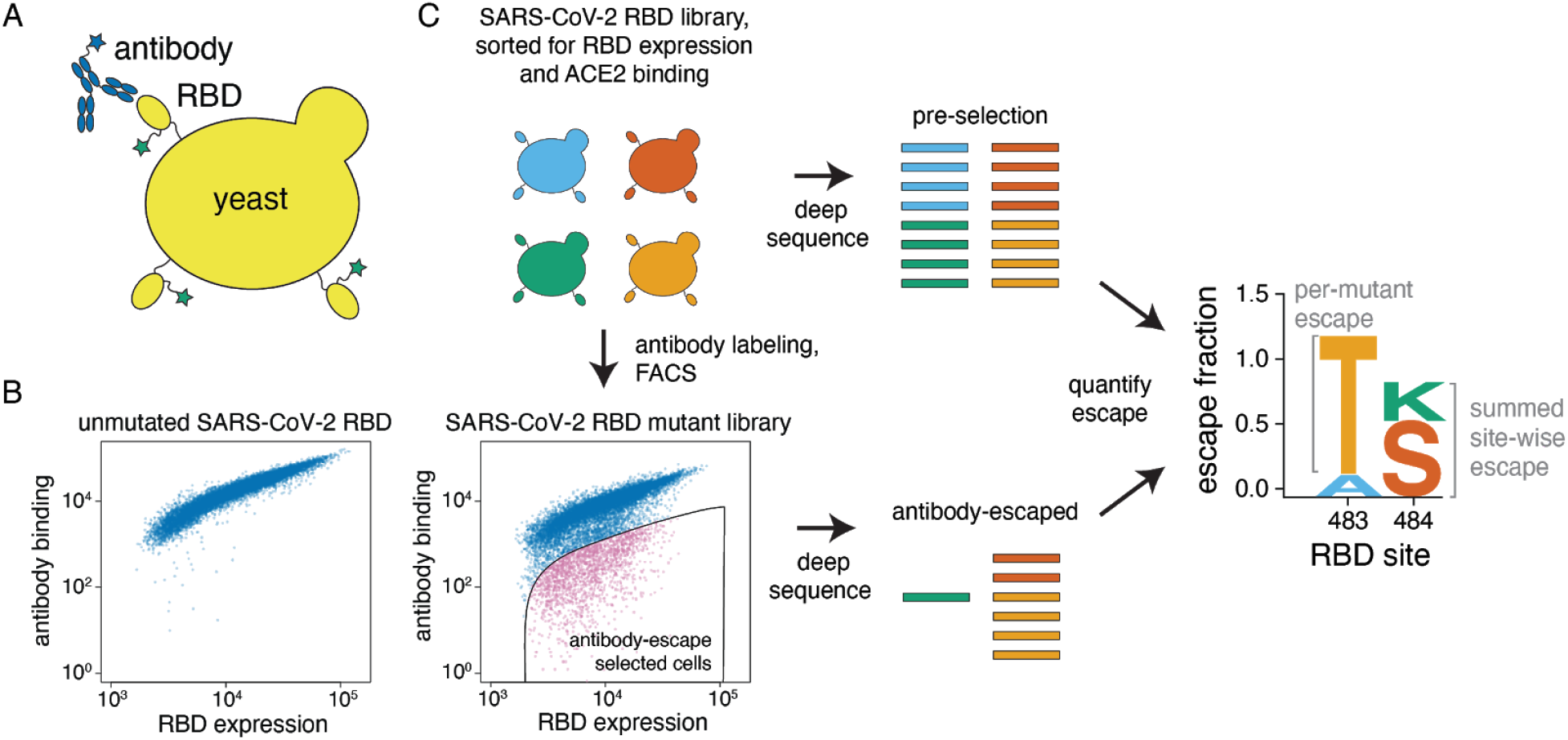
A yeast-display system to completely map SARS-CoV-2 RBD antibody escape mutations. (A) Yeast display RBD on their surface. The RBD contains a c-myc tag, enabling dual-fluorescent labeling to quantify both RBD expression and antibody binding of RBD by flow cytometry. (B) Per-cell RBD expression and antibody binding as measured by flow cytometry for yeast expressing unmutated RBD and one of the RBD mutant libraries. (C) Experimental workflow. Yeast expressing RBD mutant libraries are sorted to purge RBD mutations that abolish ACE2 binding or RBD folding. These mutant libraries are then labeled with antibody, and cells expressing RBD mutants with decreased antibody binding are enriched using FACS (the “antibody-escape” bin; see Figure S1 for gating details). Both the initial and antibody-escape populations are deep sequenced to identify mutations enriched in the antibody-escape population. The deep-sequencing counts are used to compute the “escape fraction” for each mutation, which represents the fraction of yeast cells with a given RBD mutation that falls into the antibody-escape sort bin. The escape fractions are represented in logo plots, with tall letters indicating mutations that strongly escape antibody binding.

Here, we developed a method to use these libraries to comprehensively identify mutations in the RBD that allow it to escape binding by antibodies. To eliminate RBD mutants that were completely misfolded or unable to bind ACE2, we first used fluorescence-activated cell sorting (FACS) to eliminate RBD variants with <0.01x the affinity for ACE2 compared to that of the unmutated RBD (Figure S1A,B). We reasoned this sorting would purge the libraries of completely nonfunctional RBD mutants, but retain mutants with decreased ACE2 affinity that might enable antibody escape (Baum et al., 2020a; Rockx et al., 2010). We then incubated the ACE2-sorted yeast libraries with an anti-RBD antibody (see next section) and sorted for cells that expressed RBD mutants that bound substantially less antibody than unmutated SARS-CoV-2 RBD (Figure 1C, Figure S1C). We deep-sequenced the nucleotide barcodes to quantify RBD variant frequencies in the initial ACE2+ population and the antibody-escape population (Figure 1C). We quantified the effect of each RBD mutation by estimating the fraction of cells expressing that mutation that fell into the antibody-escape sort bin, and termed this quantity the mutation’s “escape fraction”. We represented the escape fractions using logo plots (Figure 1C).

### Mapping escape from each of 10 human monoclonal antibodies

We applied our escape-mutation mapping to 10 human monoclonal antibodies: 9 neutralizing antibodies isolated from SARS-CoV-2 convalescent patients (Zost et al., 2020b), and a recombinant form of one cross-reactive non-neutralizing antibody isolated from a convalescent SARS-CoV-1 patient (rCR3022) (Huo et al., 2020; ter Meulen et al., 2006; Tian et al., 2020; Yuan et al., 2020). All 10 antibodies bind the SARS-CoV-2 RBD with high affinity, but they differ in their neutralization potencies, extent to which they compete with ACE2 for RBD binding, and cross-reactivity with SARS-CoV-1 (Figure 2A) (Yuan et al., 2020; Zost et al., 2020a).

**Figure 2.**
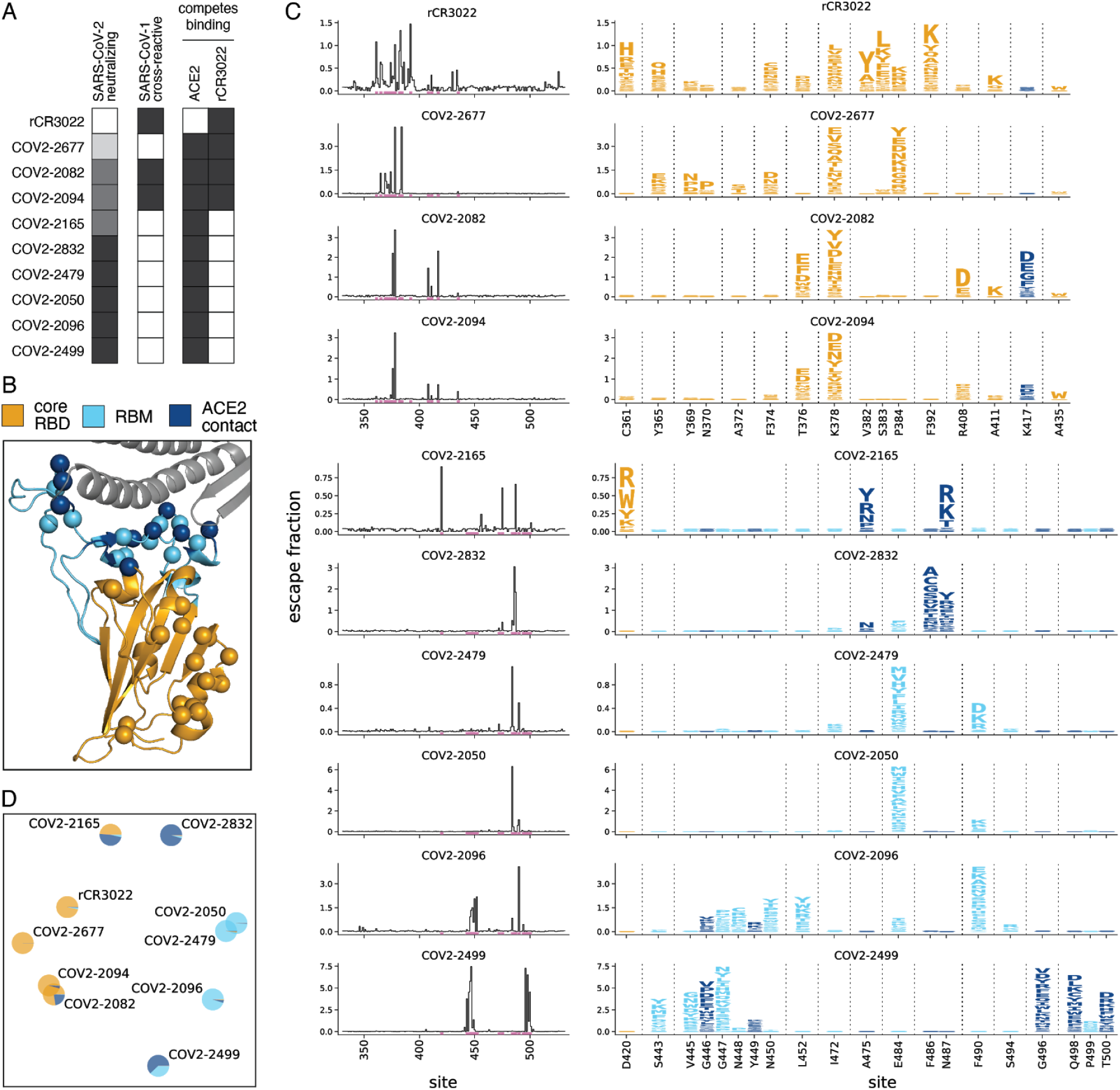
Complete maps of escape mutations from 10 human monoclonal antibodies. (A) Properties of the antibodies as reported by Zost *et al*. (2020a). SARS-CoV-2 neutralization potency is represented as a gradient from black (most potent) to white (non-neutralizing). Antibodies that bind SARS-CoV-1 spike or compete with RBD binding to ACE2 or rCR3022 are indicated in black. (B) Structure of the SARS-CoV-2 RBD (PDB: 6M0J, (Lan et al., 2020)) with residues colored by whether they are in the core RBD distal from ACE2 (orange), in the receptor-binding motif (RBM, light blue), or directly contact ACE2 (dark blue). ACE2 is in gray. RBD sites where any antibody in the panel selects escape mutations are indicated with spheres at their alpha carbons. (C) Maps of escape mutations from each antibody. The line plots show the total escape at each RBD site (sum of escape fractions of all mutations at that site). Sites with strong escape mutations (indicated by purple at bottom of the line plots) are shown in the logo plots. Sites in the logo plots are colored by RBD region as in (B), with the height of each letter representing the escape fraction for that mutation. Note that different sites are shown for the rCR3022-competing antibodies (top four) and all other antibodies (bottom six). (D) Multidimensional scaling projection of the escape-mutant maps, with antibodies having similar escape mutations drawn close together. Each antibody is shown with a pie chart that uses the color scale in (B) to indicate the RBD regions where it selects escape mutations.

We mapped escape mutations for each of the 10 antibodies in biological duplicate by applying the workflow in Figure 1C to each of our two independently generated RBD mutant libraries (Figures S1C, S2). We determined the effect of each mutation on antibody escape (the “escape fraction”, Figure 1C) after applying quality-control filters to remove RBD mutants with low expression, ACE2 binding, or sequencing counts (see Methods for details). The resulting escape fraction measurements correlated strongly between the duplicate mutant libraries (Figure S2), and for the rest of this paper we report the average measurements across libraries. Note that the magnitude of the measured effects of mutations on antibody escape depends on the antibody concentration and the flow cytometry gates applied (Figure S1C), meaning that the escape fractions are comparable across sites for any given antibody, but are not precisely comparable among antibodies without external calibration.

The effects of mutations on antibody escape are summarized in Figure 2C (see Supplemental File 1 for raw data). Each antibody is escaped by mutations at just a small subset of residues in the RBD. In general, rCR3022 and the three antibodies that compete with rCR3022 for RBD binding are escaped by mutations in the core RBD distal from the ACE2 receptor binding motif (RBM) (Figure 2A,B,C). The remaining antibodies are escaped primarily by mutations in the RBM of the RBD, including at ACE2 contact residues (Figure 2C). Notably, the escape mutations for the most potently neutralizing antibodies fall mostly in the RBM (Figure 2A,C), consistent with prior studies showing that potent anti-RBD neutralizing antibodies often strongly compete with ACE2 binding (Brouwer et al., 2020; Cao et al., 2020; Huang et al., 2020; Ju et al., 2020; Liu et al., 2020; Rogers et al., 2020; Seydoux et al., 2020; Wec et al., 2020; Wu et al., 2020; Zost et al., 2020a).

However, the escape-mutation maps are far more nuanced than can be represented by simply grouping the RBD into broad antigenic regions. While a few antibodies have extremely similar escape mutations (*e*.*g*., COV2-2082 is similar to COV2-2094, and COV2-2479 is similar to COV2-2050), antibodies that target the same broad region of the RBD often have distinct escape mutations (*e*.*g*., COV2-2832 and COV2-2499 are escaped by entirely non-overlapping sets of escape mutations in the RBM). There is also heterogeneity in which specific amino acid mutations mediate escape. At some selected sites, many mutations confer escape (*e*.*g*., site 378 for COV2-2677 or site 490 for COV2-2096). But at other sites, only certain mutations confer escape: for instance, only negatively-charged amino acids at site 408 escape COV2-2082, and only mutations at site 372 that introduce a serine or threonine (creating an N-linked glycosylation motif at site 370) escape COV2-2677.

To better compare the escape maps across antibodies, we used multidimensional scaling to project the similarity in escape mutations into a two-dimensional plot (Figure 2D). In this plot, the distance between antibodies increases as their escape mutations become more distinct. In addition, the pie chart colors in Figure 2D indicate the regions of the RBD where mutations confer escape. This plot makes clear that antibodies that target similar regions of the RBD sometimes but not always have similar escape mutations: for instance, COV2-2479, COV2-2050, and COV2-2096 all target the RBM–but only the first two of these antibodies cluster closely in Figure 2D. Overall, the two-dimensional projection in Figure 2D provides an intuitive way to visualize the relationships among antibodies in the space of immune-escape mutations, similar to how dimensionality reduction techniques such as t-SNE or UMAP help visualize high-dimensional single-cell transcriptomic data (Amir et al., 2013; Becht et al., 2019).

To independently validate the escape maps, we tested key escape mutations in neutralization assays using spike-pseudotyped lentiviral particles (Crawford et al., 2020a). The agreement between the escape maps and neutralization assays was excellent (Figure 3, Figure S3A) and validated the subtle differences between antibodies. For instance, as indicated by the maps, a mutation at site 487 escapes both COV2-2165 and COV2-2832, but a mutation at site 486 only escapes COV2-2832 (Figure 3). We also validated the map for the non-neutralizing antibody rCR3022 by showing that mutations had the expected effects on binding of this antibody to mammalian-cell expressed RBD (Figure S3B-E).

**Figure 3.**
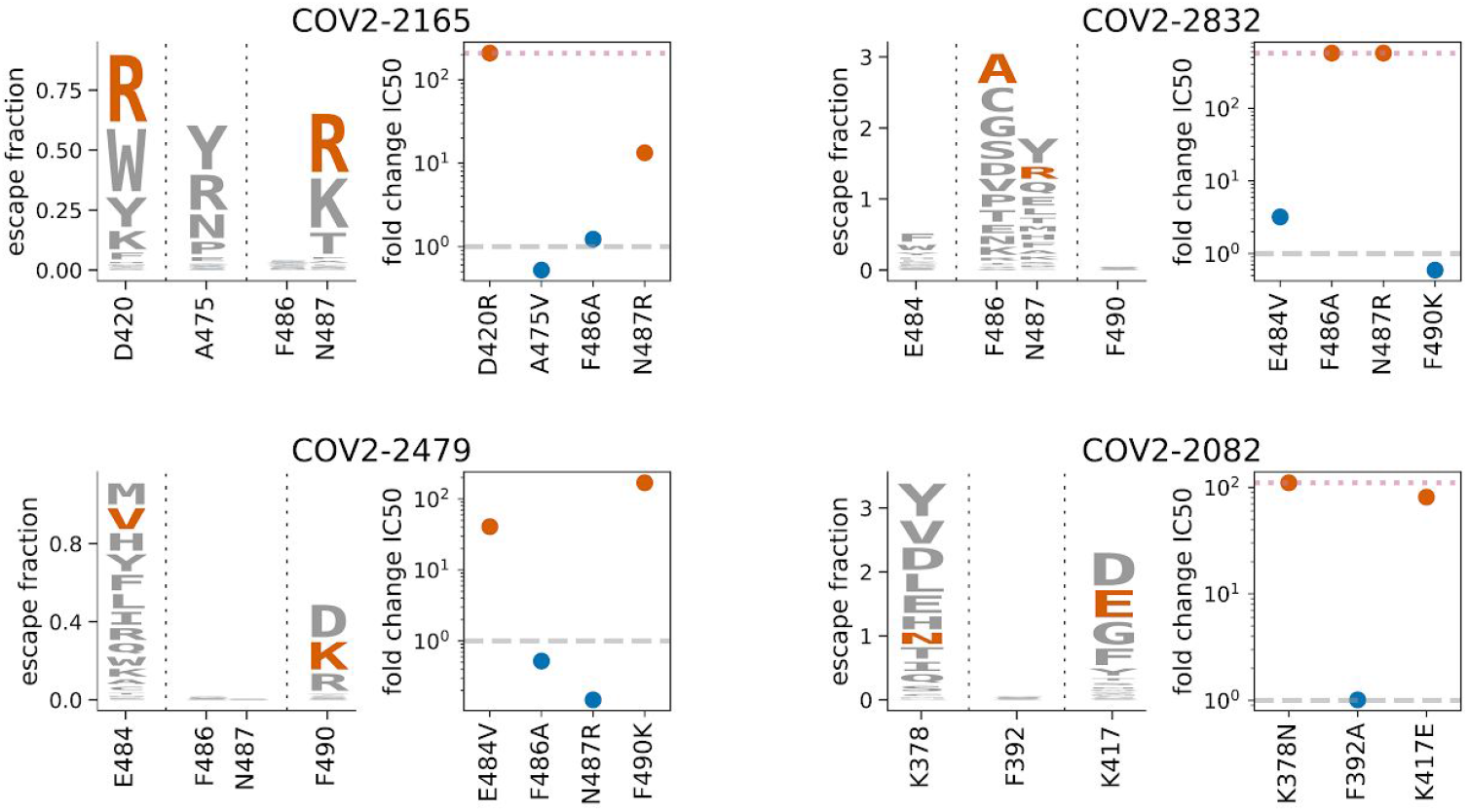
Neutralization assays validate antibody escape maps. For each of the four indicated antibodies, we chose two mutations that our maps indicated should escape antibody binding, and one or two mutations that should not escape binding. Logo plots show the escape maps for the sites of interest, with the tested mutations that should escape antibody binding in red. Dot plots show the fold change in neutralization (inhibitory concentration 50%, IC50) relative to the unmutated (wildtype) spike measured using spike-pseudotyped lentiviral particles. Fold changes greater than one (dashed gray line) mean a mutation escapes antibody neutralization. Points in red correspond to the mutations expected to mediate escape, and those in blue correspond to mutations not expected to escape (blue letters are not visible in the logo plots as they do not have substantial effects in the mapping). The dotted pink line at the top of some plots indicates the upper limit to the dynamic range; points on the line indicate a fold change greater than or equal to this value. See Figure S3A for the raw neutralization curves, and Figures S3B,C for similar validation for the non-neutralizing antibody rCR3022.

### Structural data partially but not completely explain the escape maps

We next examined the extent to which the escape maps could be rationalized in terms of the three-dimensional structures of the antibody-RBD complexes. We used negative-stain electron microscopy (EM) to obtain structures of five of the antibodies in complex with the RBD, and analyzed an existing structure of rCR3022 bound to RBD (Yuan et al., 2020). To enable structural interpretation of the escape mutations, we juxtaposed these structures of antibody-bound RBD with structural projections of our escape maps (Figure 4). We also created interactive structure-based visualizations of the escape maps using `dms-view` (Hilton et al., 2020) that are available at https://jbloomlab.github.io/SARS-CoV-2-RBD_MAP_Crowe_antibodies/ and enable facile analysis of escape mutations on the RBD structure.

**Figure 4.**
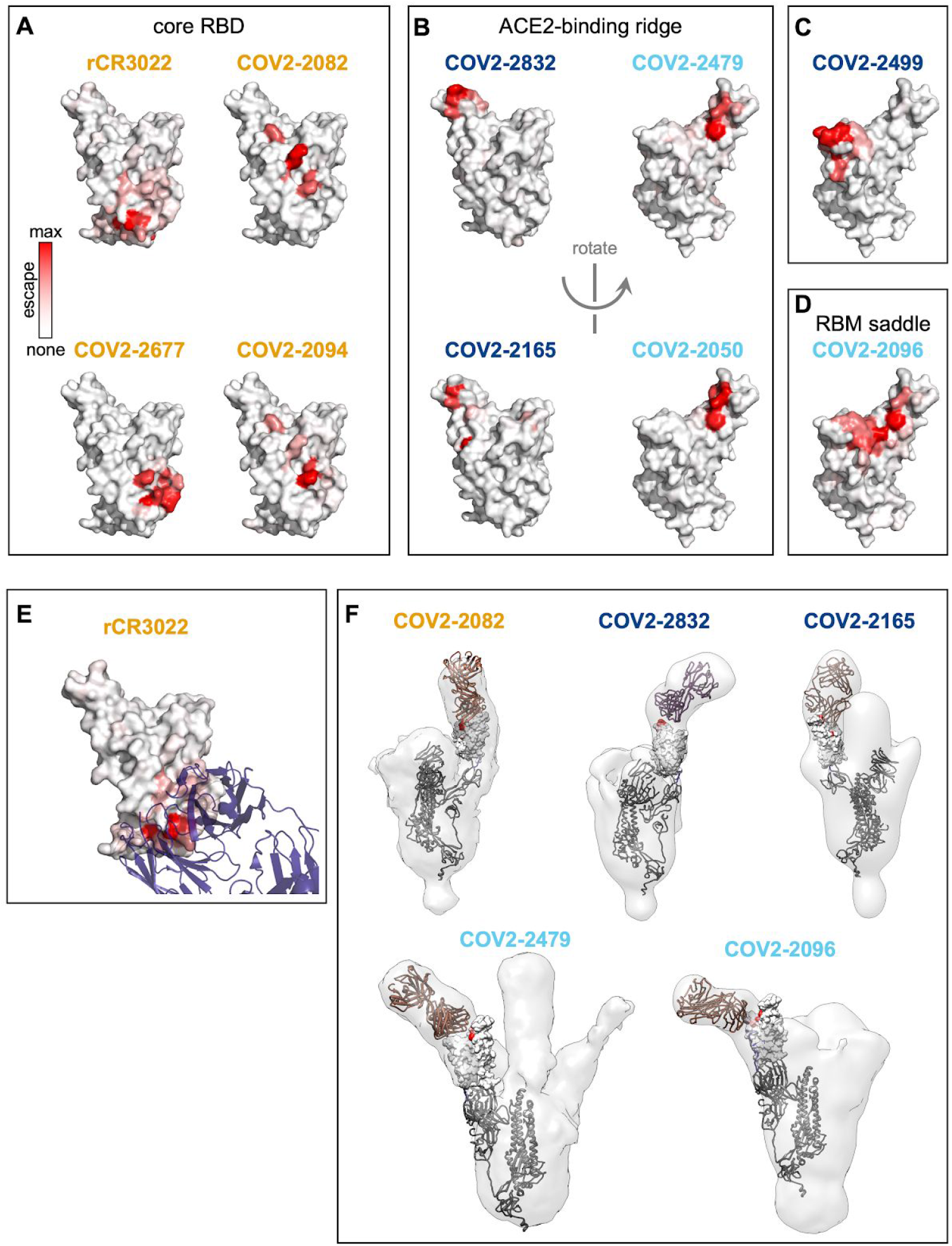
Structural mapping of antibody binding and escape. (A-D) For each antibody, the structure shows the RBD surface (PDB 6M0J) colored by the largest-effect escape mutation at each site, with white indicating no escape and red indicating the strongest escape mutation for that antibody. Antibodies are arranged so that those with similar structural epitopes are in the same panel, namely by whether their epitopes are in (A) the core of the RBD, (B) the ACE2-binding ridge, (C) the opposite edge of the RBM, or (D) the saddle of the RBM surface. (E) Crystal structure of the rCR3022-bound RBD (PDB 6W41), with Fab in purple and RBD colored according to sites of escape as in (A). (F) For 5 monoclonal antibodies, Fab bound to SARS-CoV-2 spike ectodomain trimer was visualized by negative-stain electron microscopy (EM). The RBD is modeled as a surface representation, colored according to sites of escape as in (A). Fab chains are modeled in gold. Detailed EM collection statistics are in Table S1. Antibody names are colored according to Figure 2B: core-binding, orange; RBM-binding, cyan; ACE2 contact site-binding, dark blue. See https://jbloomlab.github.io/SARS-CoV-2-RBD_MAP_Crowe_antibodies/ for interactive versions of the escape-colored structures in (A-D).

Both the antibody-RBD structures and the escape maps highlight several antigenic regions on the RBD (Figure 4). The first region, targeted by four antibodies, is on the internal face of the core RBD (Figure 4A), which is only accessible in the context of full spike protein when the RBD transitions into the “open” conformation to engage ACE2 (Huo et al., 2020; Walls et al., 2020b; Wrapp et al., 2020; Yuan et al., 2020). The remaining antibodies target several distinct regions on the RBM: four antibodies are escaped by mutations on the internal or external face of one lateral edge of the RBM (the “ACE2-binding ridge”, Figure 4B), one antibody is escaped by mutations on the external face at the opposite edge of the RBM (Figure 4C), and one antibody is escaped by mutations that bridge the exterior surface of the central concave “saddle” of the RBM (Figure 4D). In all cases, the escape mutations fall in or near the structurally defined contact surface between the antibody and RBD (Figure 4E,F). In some cases, the negative-stain EM explains specific features of the escape-mutant maps (Figure 4F). For instance, COV2-2165 is strongly escaped by mutations at site D420 in addition to the ACE2-binding ridge, suggesting a binding footprint that extends beyond the ACE2-binding ridge. This hypothesis is supported by negative-stain EM data, which shows differences in the binding approach of COV2-2165 relative to that of COV2-2832, another ACE2-binding ridge antibody that is not escaped by mutations at D420 (Figure 4F).

However, the escape-mutation maps contain substantial information beyond what can be gleaned from structure alone. For example, COV2-2832 and COV2-2479 both target the ACE2-binding ridge, but have non-overlapping escape mutations on different faces of the ridge (Figure 2C, Figure 4). Similarly, while the negative-stain EM structures show that COV2-2165 and COV2-2832 both bind the ACE2-binding ridge, and the two antibodies select escape mutations very close to one another in the three-dimensional structure (Figure 4B, left, Figure 4F), there are important differences. For instance, COV2-2832 is escaped by mutations at sites F486 and N487, while COV2-2165 is only escaped by mutations at site N487 (Figure 2, Figure 4; validated by neutralization assays in Figure 3). In addition, while some antibodies (e.g., COV2-2096) can be escaped by mutations across a wide swath of the RBD surface, others (*e*.*g*., COV-2050) are only sensitive to mutations at a handful of sites.

The fact that the escape mutations occur at only a subset of sites in the antibody-RBD interfaces is consistent with classical biochemical studies showing that protein-protein binding interfaces can be dominated by “hot spots” that contribute most of the binding energy (Clackson and Wells, 1995; Cunningham and Wells, 1993), and more recent work showing that the functional and structural epitopes of anti-viral antibodies are often distinct (Dingens et al., 2019). From a therapeutic standpoint, these results emphasize the value of directly mapping escape mutations when considering the potential for viral antibody escape. For instance, our results suggest that it should be possible to make effective cocktails of antibodies with similar structural epitopes but orthogonal escape mutations, such as COV2-2165 + COV2-2479 or COV2-2499 + COV2-2050.

### Functional and evolutionary constraint on antibody-escape mutations

Our complete maps of escape mutations enable us to assess the potential for SARS-CoV-2 to evolve to escape antibodies targeting the RBD. We first examined whether the antibody-escape mutations identified in our study are present in viruses circulating in the human population. Of 93,858 SARS-CoV-2 sequences in the GISAID database as of September 6, 2020, there were 5 or more naturally occurring mutants at 14 of the 36 RBD sites where mutations escape at least one antibody (Figure 5A, Figure S4). However, mutations at all these sites are present only at very low frequency (<0.1% of viral sequences). The antibody-escape sites with naturally occurring mutations include sites 484 and 490, where other studies have recently reported selecting mutations that escape monoclonal antibodies or sera containing polyclonal antibodies (Baum et al., 2020a; Li et al., 2020; Weisblum et al., 2020). Overall, these results show that while the vast majority of viruses remain susceptible to all antibodies examined here, there is nascent low-level viral genetic variation at some key sites of escape mutations.

**Figure 5.**
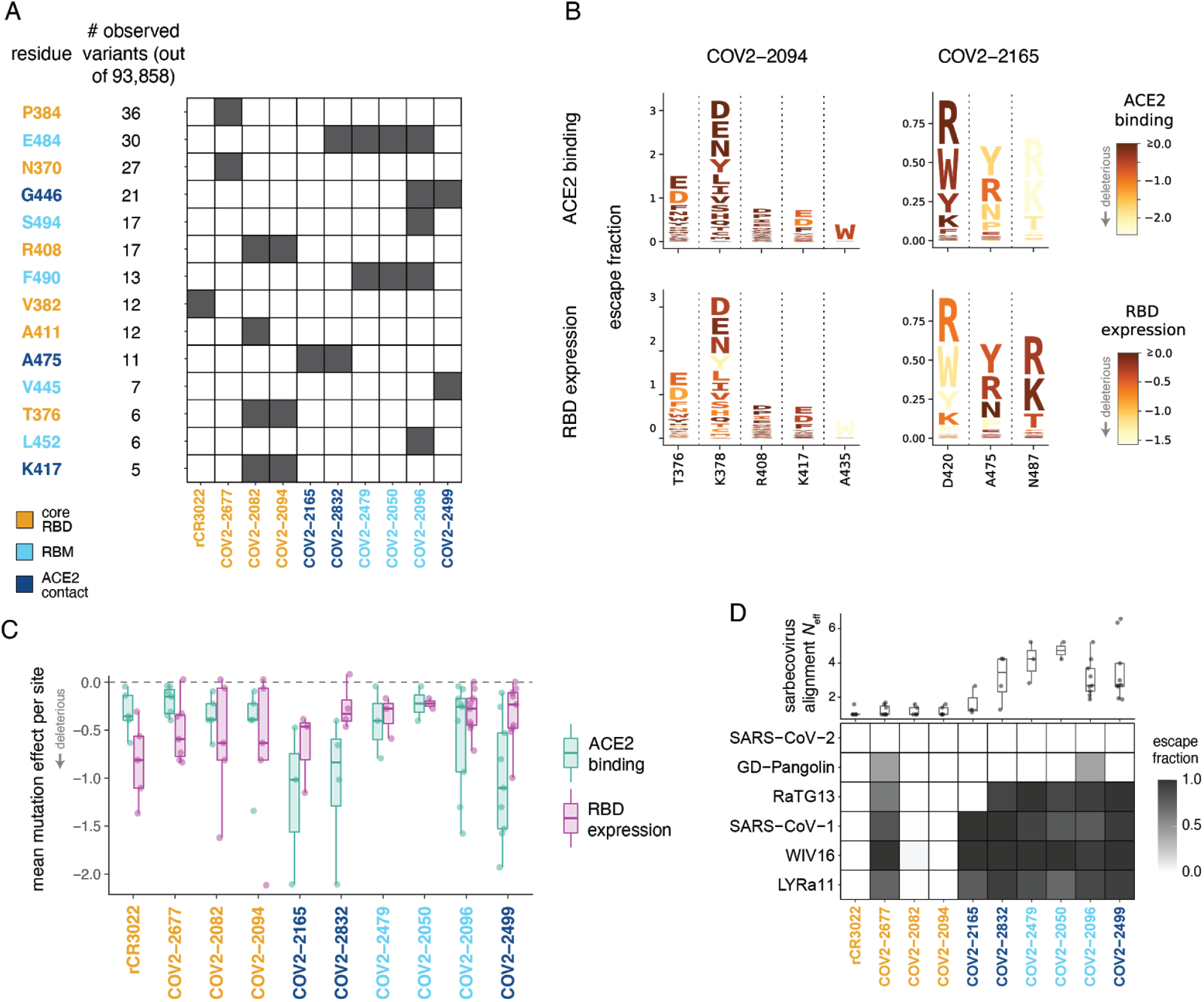
Functional and evolutionary constraint on antibody escape mutations. (A) Variation at sites of antibody escape among currently circulating SARS-CoV-2 viruses. For each site of escape from at least one antibody, we counted the sequences in GISAID with an amino-acid change (there were 93,858 sequences at the time of the analysis). Sites with at least 5 GISAID variants are shown ordered by count; Black cells indicate antibodies with escape mutations at that site. Sites are in orange for the core RBD, light blue for the RBM, and dark blue for ACE2 contact residues. Antibodies are colored according to where the majority of their sites of escape fall. Figure S4 shows similar data broken down by amino-acid change and without count thresholding. (B) Escape maps (as in Figure 2C), with letters colored according to how deleterious mutations are for ACE2 binding or RBD expression effects (Starr et al., 2020). Only sites of escape for each antibody are depicted. Similar logo plots for all antibodies are shown in Figure S5. (C) Mutational constraint on sites of escape. For each antibody, the mean effects of all 19 possible amino acid mutations at sites of escape on ACE2 binding and RBD expression are shown. (D) Top: effective number of amino acids (*N*_eff_) in the sarbecovirus RBD alignment at sites of escape for each antibody. *N*_eff_ is a measure of the variability of a site (the exponentiated Shannon entropy), and ranges from 1 for a position that is conserved across all sequences to an upper limit of 20 for a site where all amino acids are present at equal frequency. Bottom: escape fraction for each sarbecovirus RBD homolog from the yeast display selections.

To better assess the potential for future viral genetic variation, we quantified the functional constraint on sites of escape using existing deep mutational scanning measurements of how RBD mutations affect ACE2-binding and expression of properly folded RBD protein (Starr et al., 2020). Figure 5B shows the escape maps for two antibodies colored by the functional effects of mutations (comparable data for all antibodies are in Figure S5). It is obvious from Figure 5B that some escape mutations from the core-RBD-directed antibody COV2-2094 are deleterious for expression of properly folded RBD (*e*.*g*., mutations at site 435), whereas some escape mutations from the RBD-directed antibody COV2-2165 are deleterious for ACE2 binding (*e*.*g*., mutations at site 487). To quantify this trend, we determined the mean functional effect of all mutations at each site of escape from each antibody (Figure 5C). At a broad level, sites of escape from antibodies targeting the RBM and especially ACE2-contact residues are often constrained by how mutations affect ACE2 binding. On the other hand, sites of escape from antibodies targeting the core RBD are often constrained by how mutations affect RBD folding and expression (Figure 5B). These observations highlight how some antibodies target RBD sites that are functionally constrained and thus may have reduced potential for evolution.

We also examined the ability of each antibody to bind RBDs from other SARS-related coronaviruses (sarbecoviruses). To do this, we included in our libraries the unmutated RBDs from two close relatives of SARS-CoV-2 (RaTG13 (Zhou et al., 2020b) and GD-Pangolin (Lam et al., 2020)), along with SARS-CoV-1 and two of its close relatives (WIV16 (Yang et al., 2016) and LYRa11 (He et al., 2014)). Using the same approach employed to measure the effects of mutations to SARS-CoV-2, we quantified the ability of each antibody to bind these RBD homologs. We found a stark difference in cross-sarbecovirus reactivity between antibodies targeting the core RBD and those targeting the RBM (Figure 5D). Three of four antibodies targeting the core RBD bound to all five RBD homologs, whereas RBM-directed antibodies only bound the two homologs most closely related to SARS-CoV-2 (GD-Pangolin and RaTG13). This pattern is explained by the evolutionary conservation at sites of escape (Figure 5D, top): in general, sites of escape from antibodies targeting the RBD core are mostly conserved across sarbecoviruses, while sites of escape from RBM-directed antibodies are highly variable across sarbecoviruses. The only exception is COV2-2677, which does not bind any other RBD homologs despite targeting conserved sites in the core RBD: this discrepancy is explained by the A372T escape mutation, which restores an N370 glycosylation motif that is present in all sarbecoviruses except SARS-CoV-2. These results show that antibodies targeting the conserved core RBD are more likely than antibodies targeting the RBM to provide pan-sarbecovirus immunity.

### Escape maps predict results of antibody selection experiments and inform design of cocktails

We next examined if the escape maps accurately predicted the mutants selected when virus is grown in the presence of antibody. To investigate this, we used a recombinant replication-competent vesicular stomatitis virus (VSV) expressing the SARS-CoV-2 spike in place of the endogenous VSV glycoprotein (G) (Case et al., 2020). Such viruses provide a facile system to select for spike mutations that evade antibody neutralization (Case et al., 2020; Dieterle et al., 2020; Weisblum et al., 2020). We chose five potently neutralizing antibodies (IC50 values ranged from 15 to 150 ng/mL), and used a high-throughput quantitative real-time cell analysis assay (Gilchuk et al., 2020a, 2020b) to select viral mutants that could escape each individual antibody at a concentration of 5 μg/mL, performing between 16 and 56 individual replicates for each antibody (Figure 6A and S6A,B,C). For four of the five antibodies, this process selected viral variants that we confirmed resisted neutralization by 10 μg/mL of the antibody used for the selection (Figure 6A). For one antibody (COV2-2165), no escape mutants were detected even in 56 attempted replicates (Figure 6A). We sequenced the antibody-selected escape viruses, and in all cases they carried RBD mutations that the escape maps indicated mediate strong escape (examine the mutations in Figure 6A on the maps in Figure 2C).

**Figure 6.**
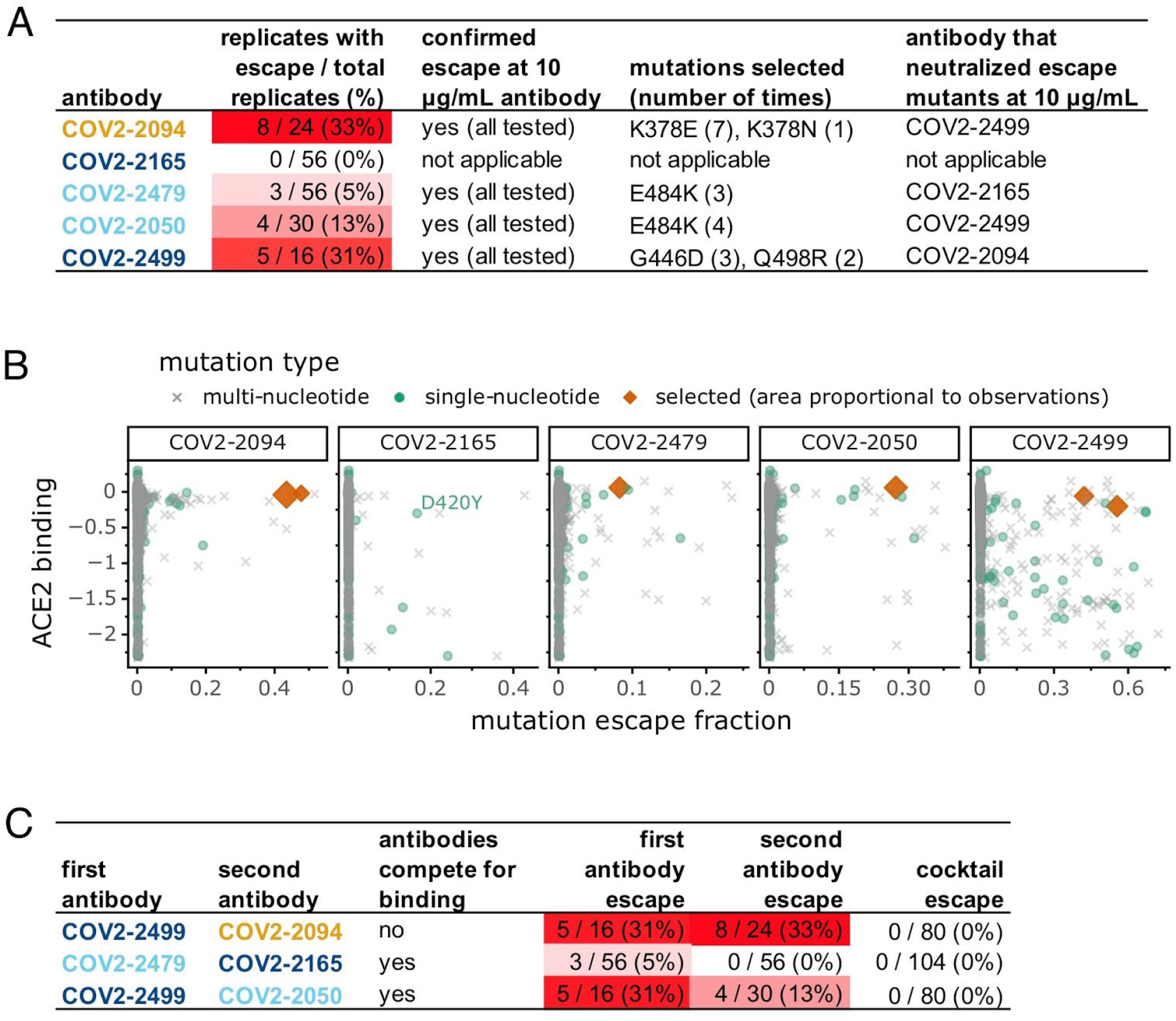
Viral escape-mutant selections with individual antibodies and antibody cocktails. (A) Results of viral selections with five individual monoclonal antibodies. The number of replicates where escape variants were selected are indicated, color coded according to whether escape was selected frequently (red) or rarely (white). The mutations present in the RBD of the selected escape variants are indicated. (B) Each point represents a different amino-acid mutation to the RBD, with the x-axis indicating how strongly the mutation ablates antibody binding in our escape maps (larger values indicate more escape from binding) and the y-axis indicating how the mutation affects ACE2 binding (negative values indicate impaired ACE2 binding). The point shapes indicate whether or not mutations are accessible by single-nucleotide changes, and whether they were selected in viral escape experiments. All selected mutations were accessible by single-nucleotide changes. Note that the only accessible escape mutation from COV2-2165 that is not deleterious to ACE2 binding is D420Y, but this mutation is highly deleterious to expression of properly folded RBD (Figure 5B and S5). (C) Results of viral selections with antibody cocktails, with the last three columns showing the number of replicates with escape out of the total tested. The data for the single antibodies are repeated from (A). In all panels, antibody names are colored according to where in the RBD the majority of their sites of escape fall: orange for the core RBD, light blue for the RBM, and dark blue for ACE2 contact residues. See Figure S6 for additional data relevant to this figure.

We next sought to understand why the antibodies selected the viral mutations that they did–and why it was not possible to select any viral mutants that escaped one of the antibodies. To do this, we considered two additional factors: which mutations are tolerated for protein function, and which mutations are accessible by single-nucleotide changes. We assessed how well mutations are tolerated functionally using deep mutational scanning measurements of how all RBD mutations affect ACE2 binding (Starr et al., 2020). We plotted all mutations in scatter plots to examine their impact on antibody escape and ACE2 binding, further stratifying by whether mutations were accessible by single-nucleotide changes to the spike gene encoded in the VSV (Figure 6B). The mutations selected by the antibodies were consistently among the ones with the largest effects on antibody escape that also did not greatly impair ACE2 binding and were accessible by single-nucleotide changes (red diamonds in Figure 6B). The antibody for which we could not select any viral escape mutants (COV2-2165) only had a single escape mutation (D420Y) that was accessible by a single-nucleotide change and not highly deleterious for ACE2 binding. However, D420Y is extremely deleterious for expression of properly folded RBD protein (Figure 5B and S5), explaining why it was not possible to select any viral escape mutants from COV2-2165. Therefore, the escape maps can be combined with deep mutational scanning of functional constraint and basic knowledge of the genetic code to predict which viral mutations are likely to arise under antibody pressure–and to identify antibodies for which escape mutations are unlikely.

One approach to thwart the risk of viral escape that is inherent in monotherapy approaches is to use antibody cocktails (Julg et al., 2017; Wec et al., 2019). In the context of SARS-CoV-2, recent work has demonstrated that cocktails of two antibodies that do not compete for binding to the same region of spike may offer higher resistance to escape mutations (Baum et al., 2020a) while protecting animals from SARS-CoV-2 challenge (Baum et al., 2020b; Zost et al., 2020a). We hypothesized that we could leverage our escape maps to rationally design more nuanced cocktails of antibodies with distinct escape mutations, even if the antibodies recognize overlapping antigenic regions and compete for binding to spike.

We created three different two-antibody cocktails: one “conventional” cocktail of antibodies that did not compete for binding to spike protein (COV2-2499 + COV2-2094), and two cocktails of antibodies that competed for binding to the RBM region of the spike protein RBD but that our maps indicated were escaped by distinct mutations (COV2-2479 + COV2-2165 and COV2-2499 + COV2-2050) (Figure 6C and S6D). Each cocktail contained a 1:1 mix of the two constituent antibodies at a total concentration of 5 μg/mL of antibody, so that the total antibody concentration was the same as in the single-antibody selections described above. We performed between 80 and 104 escape-selection replicates with each cocktail. No cocktail escape mutants were identified in any of these replicates, despite the fact that two of the cocktails were composed of antibodies for which substantial numbers of escape mutants were selected by the individual antibodies (Figure 6C and S6C). The lack of cocktail escape mutants is likely due to the “orthogonality” of the escape mutations for the individual antibodies, as viruses with the mutations selected by each single antibody were sensitive to the other antibody in the cocktail (Figure 6A). Overall, these results demonstrate how complete escape maps can inform the design of “non-conventional” cocktails of antibodies that compete for binding to the antigen but are nonetheless resistant to viral escape because they have orthogonal escape mutations (*e*.*g*., the cocktail COV2-2499 + COV2-2050).

## Discussion

We have described an approach to completely map mutations to the SARS-CoV-2 RBD that escape antibody binding. Unlike traditional selection experiments that only identify a handful of the possible escape mutations, our method completely maps mutations that escape antibody binding. These maps complement structure-based approaches that define the physical interface between an antibody and virus but do not directly measure how mutations affect antibody binding.

The escape maps reveal remarkable nuance in which mutations escape individual antibodies. Our maps and corroborating structural data show that at a superficial level, the antibodies target just a few patches on the surface of the RBD that likely correspond to “antigenic regions” that have been defined using other approaches (Barnes et al., 2020b; Brouwer et al., 2020; Liu et al., 2020; Robbiani et al., 2020; Rogers et al., 2020; Wec et al., 2020; Zost et al., 2020a). However, the fine details of the escape maps show that the effects of specific mutations can vary dramatically even among antibodies that superficially target the same region. For instance, antibodies COV2-2479, COV2-2050, and COV2-2832 all target the RBD ACE2-binding ridge–but while the first two have nearly identical escape mutations, the escape mutations for COV2-2832 are almost completely distinct. We speculate that these differences arise from the fact that even antibodies that physically contact a large surface area on the RBD are often only escaped by mutations at a few residues, a vivid illustration of the classically defined importance of “hot spots” in antibody-antigen binding (Bogan and Thorn, 1998; Dall’Acqua et al., 1998; Jin et al., 1992).

We also overlaid the escape maps with existing deep mutational scanning data on the functional consequences of mutations for the expression of properly folded RBD and its affinity for ACE2 (Starr et al., 2020). In general, the sites of escape from antibodies directed to the core RBD are constrained with respect to their effects on expression of properly folded RBD, whereas sites of escape from antibodies directed to the RBD’s receptor-binding motif are more constrained with respect to their effects on ACE2 binding. While these analyses come with the caveat that no experimental measures of the effects of mutations fully capture how they affect true viral fitness, it is nonetheless informative to assess how mutations that escape antibody binding impact the key biochemical functions of the RBD.

Remarkably, combining the escape maps with these functional measurements predicts which mutations are selected when spike-expressing virus is grown in the presence of individual antibodies. The selected viral escape mutations are consistently those that have large effects on antibody escape but little negative impact on ACE2 binding and RBD folding, and are also accessible by single-nucleotide mutations. Furthermore, one of the antibodies was highly resistant to viral escape–and we showed this could be explained by the fact that the virus has no escape mutations from this antibody that are both tolerable for RBD function and accessible by single-nucleotide changes. Therefore, complete measurements of both the antigenic and functional consequences of viral mutations provide the phenotypic data necessary to assess both the likelihood of viral escape under antibody pressure and the specific mutations that arise when escape occurs.

One immediate implication of our results is that counter to prevailing wisdom, antibody cocktails do not have to target distinct regions of the RBD in order to resist viral escape. Simple inspection of the escape maps reveals pairs of antibodies targeting the RBD’s ACE2-binding interface that share no common escape mutations, and so could be good candidates for therapeutic cocktails. Indeed, we combined our escape maps with selections on spike-expressing viruses to show that cocktails of antibodies that compete for binding to spike but have different escape mutations still resist viral escape. It is possible that such cocktails could even be preferable to cocktails of antibodies targeting distinct regions (Schmidt et al., 2015; Schommers et al., 2020), since acquiring multiple different escape mutations in the ACE2 binding interface could impose an intolerable loss of receptor binding on the virus.

Our results are also of utility for assessing if ongoing viral evolution is likely to be of antigenic consequence. The escape maps enable immediate assessment of whether mutations to the RBD alter antigenicity. At over a dozen of the sites of escape that we mapped for these antibodies, there is already low-level genetic variation among circulating SARS-CoV-2 strains. Furthermore, the high-throughput nature of our experimental approach should make it possible to rapidly generate similar maps for other monoclonal antibodies or polyclonal antibodies in sera, thereby providing quantitative experimental data that can be cross-referenced to mutations observed during genomic surveillance of circulating SARS-CoV-2 strains (Korber et al., 2020).

It is important to note that our approach maps how mutations affect antibody binding to yeast-displayed RBD, which comes with two caveats. First, our approach can only map escape from antibodies that target epitopes entirely within the RBD, and will not identify mutations that mediate escape by altering the relative positioning of the RBD in the context of full spike protein (Weissman et al., 2020; Zhou et al., 2020c). Second, although yeast do add N-linked glycans to the RBD at the same sites as human cells (Chen et al., 2014), these glycans are more mannose-rich (Hamilton et al., 2003), which could affect binding by antibodies with glycan-rich epitopes. However, despite these potential caveats, all the mapped escape mutations that we tested had the expected effects in the context of spike-pseudotyped lentiviral or VSV particles. In addition, our approach can map mutations that escape binding by non-neutralizing as well as neutralizing antibodies, and we successfully validated mutations that ablated binding by a non-neutralizing antibody using mammalian-cell produced RBD.

Some viruses, such as measles, are antigenically stable such that immunity from an initial infection or vaccination typically provides life-long protection (Linnemann, 1973; Panum, 1939). Others, such as influenza virus, undergo rapid antigenic drift, such that immunity elicited against one viral strain can be ineffective against that strain’s descendents just a few years later (Lee et al., 2019; Smith et al., 2004). The extent to which mutations that substantially affect the antigenicity of SARS-CoV-2 will fix during viral evolution remains an open question. The escape-mutation maps we have generated, as well our methodology for rapidly creating such maps for additional antibodies and sera, should help answer this question by facilitating assessment of the antigenic consequences of mutations observed during viral surveillance.

## Acknowledgments

We thank Keara Malone for experimental assistance, Mike Murphy and Neil King for rCR3022 protein, and the Flow Cytometry and Genomics core facilities at the Fred Hutchinson Cancer Research Center for experimental support, especially Dolores Covarrubias, Andy Marty, and MinJae Kim. EM data collections were conducted at the Center for Structural Biology Cryo-EM Facility at Vanderbilt University. This work was supported by the NIAID / NIH (R01AI141707 and R01AI12893 to J.D.B., T32AI083203 to A.J.G., and F30AI149928 to K.H.D.C.). T.N.S. is a Washington Research Foundation Innovation Fellow at the University of Washington Institute for Protein Design and a Howard Huges Medical Institute Fellow of the Damon Runyon Cancer Research Foundation (DRG-2381-19). This work was also supported by Defense Advanced Research Projects Agency (DARPA) grants HR0011-18-2-0001 and HR0011-18-3-0001; NIH contracts 75N93019C00074 and 75N93019C00062; NIH grants U01 AI150739, R01 AI130591 and R35 HL145242, the Dolly Parton COVID-19 Research Fund at Vanderbilt, and a grant from Fast Grants, Mercatus Center, George Mason University. S.J.Z. was supported by NIH T32 AI095202. J.E.C. is a recipient of the 2019 Future Insight Prize from Merck KGaA, which supported this work with a grant. J.D.B. is an Investigator of the Howard Hughes Medical Institute. The content is solely the responsibility of the authors and does not necessarily represent the official views of the US government or the other sponsors.

## Author contributions

Conceptualization, A.J.G., T.N.S., S.Z., J.E.C., and J.D.B.; Methodology, A.J.G., T.N.S., S.Z., E.B., P.G., R.S.N., R.E.S., N.S., P.W.R., Z.L., S.P.J.W, R.H.C, and J.D.B.; Investigation, A.J.G., T.N.S., S.Z., P.G., and E.B.; Code, A.J.G., T.N.S., and J.D.B.; Formal Analysis, A.J.G., T.N.S., and J.D.B.; Validation, A.J.G., A.N.L., R.E., and K.H.D.C.; Writing – Original Draft, A.J.G., T.N.S. and J.D.B.; Writing – Review and Editing, all authors; Supervision, J.E.C. and J.D.B.

## Declarations of Interests

J.E.C. has served as a consultant for Sanofi; is on the Scientific Advisory Boards of CompuVax and Meissa Vaccines; is a recipient of previous unrelated research grants from Moderna and Sanofi; and is a founder of IDBiologics. Vanderbilt University has applied for patents concerning SARS-CoV-2 antibodies analyzed in this work. S.P.J.W. and P.W.R. have filed a disclosure with Washington University for the recombinant VSV. The other authors declare no competing interests.

## Methods

### Data and code availability

We provide data and code in the following ways:

- Raw data tables of single-mutation escape fractions, averaged across libraries (Supplementary File 1, and GitHub: https://github.com/jbloomlab/SARS-CoV-2-RBD_MAP_Crowe_antibodies/blob/master/results/supp_data/MAP_paper_antibodies_raw_data.csv)
- Raw data table of single-mutation escape fractions, measurements for individual library replicates (GitHub: https://github.com/jbloomlab/SARS-CoV-2-RBD_MAP_Crowe_antibodies/blob/master/results/escape_scores/escape_fracs.csv)
- Illumina sequencing counts for each barcode in each antibody escape bin (GitHub: https://github.com/jbloomlab/SARS-CoV-2-RBD_MAP_Crowe_antibodies/blob/master/results/counts/variant_counts.csv)
- The complete computational pipeline to analyze these data (GitHub: https://github.com/jbloomlab/SARS-CoV-2-RBD_MAP_Crowe_antibodies)
- A Markdown summary of the organization of analysis steps, with links to key data files and Markdown summaries of each step in the analysis pipeline (Github: https://github.com/jbloomlab/SARS-CoV-2-RBD_MAP_Crowe_antibodies/blob/master/results/summary/summary.md)
- All raw sequencing data are uploaded to the NCBI Short Read Archive (BioProject: PRJNA639956, BioSample: SAMN16054076)
- Electron density maps for the Fab/SARS-CoV-2 S complex are available from the Electron Microscopy Data Bank under the following accession codes: EMD-22627 and EMD-22628 (see also Table S1).

### Description of RBD deep mutational scanning library

The yeast-display RBD mutant libraries are identical to those previously described (Starr et al., 2020). Briefly, mutant libraries containing an average of 2.7 amino-acid mutations per variant were constructed in the spike receptor binding domain (RBD) from SARS-CoV-2 (isolate Wuhan-Hu-1, Genbank accession number MN908947, residues N331-T531). Duplicate mutant libraries were generated, and contain 3,804 of the 3,819 possible amino-acid mutations, with >95% present as single mutants. Each RBD variant was linked to a unique 16-nucleotide barcode sequence to facilitate downstream sequencing. The RBD mutant library also contained non-mutated sarbecovirus RBD homologs, RaTG13, Genbank MN996532; GD-Pangolin consensus from Lam et al. (2020); SARS-CoV-1 Urbani, Genbank AY278741; WIV16, Genbank KT444582; and LYRa11, Genbank KF569996.

### Human monoclonal antibodies targeting SARS-CoV-2 RBD

The 9 human monoclonal antibodies isolated from SARS-CoV-2 convalescent patients were produced as described in Zost *et al*. (2020b). The recombinant CR3022 antibody (rCR3022), was kindly provided by Neil King and Mike Murphy, University of Washington, Institute for Protein Design, based on the sequence reported by ter Meulen et al. (2006). All antibodies were expressed as human IgG.

Properties of the ten antibodies represented in Figure 2A were reported by Zost et al. (2020a): SARS-CoV-2 neutralization potency (black, IC_50_ <150 ng/mL; dark gray, 150-1,000; light gray, 1,000-1:10,000; white, no detectable inhibition); SARS-CoV-1 spike binding via ELISA (black, detectable; white, no detectable binding); potency of ACE2 competition via ACE2-blocking ELISA (black, IC_50_ < 150 ng/mL; white, no competition); and rCR3022 competition via ELISA (black, <25% baseline rCR3022 binding when pre-incubating with saturating antibody; white, >60% of baseline rCR3022 binding).

### Fluorescence activated cell sorting (FACS) of yeast libraries to eliminate mutants that are completely non-folded or do not bind ACE2

Libraries were sorted for RBD expression and ACE2 binding to eliminate RBD variants that are completely misfolded or non-functional (Figure S1A,B). We chose staining and sorting conditions that would select for variants with ACE2 affinity comparable to or better than RaTG13, the homolog with the lowest affinity that still marginally mediates cell entry (Shang et al., 2020). Yeast library aliquots of 18 OD units (∼1e8 cfus) were thawed into 180 mL SD-CAA (6.7 g/L Yeast Nitrogen Base, 5.0 g/L Casamino acids recipe, 1.065 g/L MES, and 2% w/v dextrose) and grown overnight shaking at 30°C, 280rpm. 33.3 OD units were back-diluted into 50 mL SG-CAA+0.1% dextrose (SD-CAA with 2% w/v galactose and 0.1% w/v dextrose in place of 2% dextrose) to induce RBD surface expression. Yeast were induced for 16-18 h at 23°C with mild agitation. 25 OD units of cells were washed twice with PBS-BSA (1x PBS with 0.2 mg/mL BSA), and incubated with 1e-8 M biotinylated ACE2 (ACROBiosystems AC2-H82E6) for 1 h at room temperature. Cells were washed with ice-cold PBS-BSA before secondary labeling for 1 h at 4°C in 3 mL1:200 PE-conjugated streptavidin (Thermo Fisher S866) to label for bound ACE2, and 1:100 FITC-conjugated anti-Myc (Immunology Consultants Lab, CYMC-45F) to label for RBD surface expression. Labeled cells were washed twice with PBS-BSA and resuspended in 2.5 mL PBS. FACS was used to enrich RBD libraries for cells capable of binding ACE2, via a selection gate drawn to capture unmutated SARS-CoV-2 cells labeled at 1% the ACE2 concentration of the library samples (*i*.*e*., 1e-10 M ACE2) (Figure S1B). 15 million ACE2+ cells were collected for each library, grown overnight in SD-CAA medium, and stored at -80°C in 9 OD unit (∼5e7 cfus) aliquots.

### Sorting of yeast libraries to select mutants that escape binding by antibodies

Antibody selection experiments were performed in biological duplicate using the independently generated mutant RBD libraries. One 9 OD unit aliquot of each ACE2+-enriched RBD library was thawed and grown overnight in 45 mL SD-CAA. Libraries were induced as described above. Induced cultures were washed and incubated with 400 ng/mL antibody for 1 h at room temperature with gentle agitation, followed by secondary labeling with 1:100 FITC-conjugated anti-Myc to label for RBD expression and 1:200 PE-conjugated goat anti-human-IgG (Jackson ImmunoResearch 109-115-098) to label for bound antibody. A flow cytometric selection gate was drawn to capture unmutated SARS-CoV-2 cells labeled at 1% the antibody concentration of the library samples (Figure S1C). Libraries were sorted to select RBD variants that reduce antibody binding and fall into this selection gate. For each sample, approximately 10 million RBD+ cells were processed on the cytometer, with between 4e5 and 2.6e6 antibody-escaped cells collected per sample (see percentages in Figure S1C for what fraction of the library had reduced binding to each antibody). Antibody-escaped cells were grown overnight in SD-CAA to expand cells prior to plasmid extraction.

### DNA extraction and Illumina sequencing

Plasmid samples were prepared from overnight cultures of antibody-escaped and 30 OD units (1.6e8 cfus) of pre-selection yeast populations (Zymoprep Yeast Plasmid Miniprep II). The 16-nucleotide barcode sequences identifying each RBD variant were amplified by PCR and prepared for Illumina sequencing exactly as described in Starr et al. (2020). Barcodes were sequenced on an Illumina HiSeq 3500 with 50 bp single-end reads. To minimize noise from inadequate sequencing coverage, we ensured that each antibody-escape sample had at least 3x as many post-filtering sequencing counts as FACS-selected cells, and reference populations had at least 2.5e7 post-filtering sequencing counts.

### Analysis of deep sequencing data to compute antibody escape fraction for each mutation

We computed escape fractions for each mutation from the counts in the Illumina deep sequencing of the 16-nucleotide barcodes as schematized in Figure 1C. We first used the `dms_variants` package (https://jbloomlab.github.io/dms_variants/, version 0.8.2) to process the Illumina sequences into counts of each barcoded RBD variant in each condition using the barcode / RBD-variant look-up table described in Starr et al. (2020). A rendering of the code that performs this variant counting is at https://github.com/jbloomlab/SARS-CoV-2-RBD_MAP_Crowe_antibodies/blob/master/results/summary/count_variants.md.

We then computed the “escape fraction” for each barcoded variant in each antibody-selected library, which we define as *E*_*v*_ = *F* × (*n*_*v*_^*post*^/*N*_*post*_)/ *n*_*v*_^*pre*^/*N*_*pre*_)/ where *F* is the total fraction of the library that escapes antibody binding (these fractions are given as percentages in the bottom two rows of Figure S1C), *n*_*v*_^*post*^ and *n*_*v*_^*pre*^ are the counts of variant *v* in the RBD library after and before enriching for antibody-escape variants with a pseudocount of 0.5 added to all counts, and 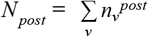 and 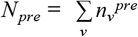 are the total counts of all variants before and after the antibody-escape enrichment. These escape fractions represent the fraction of a given variant that escape antibody binding, and should in principle range from 0 to 1. But due to statistical fluctuations in the counts sometimes the escape fractions *E*_*v*_ can be greater than one: any values of *E*_*v*_ > 1 were set to 1.

We then computationally applied two filters to remove variants that fail to express properly folded RBD and so escape antibody binding for that trivial reason rather than antibody-specific escape mutations. In principle, such variants should have been fully removed by the initial sort that only retained yeast cells with appreciable RBD expression and ACE2 binding, but in practice a small background remained as demonstrated by the fact that stop-codon variants were present at very low but still non-zero levels. For the first filter, we removed all variants with pre-selection counts lower than the counts in the 99th percentile of stop-codon-containing variant ordered by count. The logic was that this filter removed nearly all variants that were observed less frequently than stop-codon variants, which are assumed to not express properly folded RBD. For the second filter, we removed any variants that had ACE2-binding scores <-2.35 or RBD expression scores <-1.5 using the scores measured in Starr *et al*. (2020). In addition, we removed any variants that had single mutations with scores less than either of these thresholds (again using the single-mutation scores determined in Starr *et al*. (2020)) even if the variant score itself was above this threshold. The logic was that this filter removed any variants that fail to express at least low levels of properly folded ACE2. A rendering of the code that performs the computation of the escape fractions and this subsequent filtering is at https://github.com/jbloomlab/SARS-CoV-2-RBD_MAP_Crowe_antibodies/blob/master/results/summary/counts_to_scores.md.

We next deconvolved the variant-level escape fractions into escape fraction estimates for individual mutations. To do this we used global epistasis models (Otwinowski et al., 2018) as implemented in the the `dms_variants` package as detailed at (https://jbloomlab.github.io/dms_variants/dms_variants.globalepistasis.html), using the same Gaussian likelihood function as in Otwinowski *et al*. (2018). In order to make the fitting more reliable, we removed any variants with mutations not seen in at least one single-mutant variant or multiple multiple-mutant variants. We report the escape fraction on the “observed phenotype” scale: that is, we use the global epistasis models to transform the variant-level escape fractions to estimated latent phenotypes for each mutation, and then re-transform those latent phenotype estimates back through the global epistasis model. If any of these re-transformed escape fractions were not in the range between 0 and 1, they were adjusted to a minimum value of 0 or a maximum value of 1. The end result of this process was a separate estimate for each library and antibody of the escape fraction for each mutation that was not highly deleterious for expression of properly folded RBD. The correlation between these estimates for the different libraries is in Figure S2. In this paper, we report the average of the two libraries, and in the rare cases a mutation is only sampled in one library then we report the value for just that library. These values are reported in Supplementary File 1. The code that performs this global epistasis decomposition of escape scores for individual mutations is at https://github.com/jbloomlab/SARS-CoV-2-RBD_MAP_Crowe_antibodies/blob/master/results/summary/scores_to_frac_escape.md.

In some places in this paper and in Supplementary File 1, we report site-level measurements in addition to mutation-level escape scores. The first measure of site-level escape is the total site escape (total height of letter stacks, e.g. in Figure 1C), and simply represents the sum of all mutation-level escape fractions at a site. The second measure of site-level escape is the maximum escape at a site, which is just the maximum of all of the mutation-level escape fractions at the site.

### Classification of sites of escape from each antibody

For certain visualizations or analyses, it was necessary to classify which sites mediated escape from each antibody. To do this, for each antibody we identified those sites where the total site escape was >10x the median across all sites, and was also at least 10% of the maximum total site escape for any site for that antibody. We found that this heuristic reliably separated sites of clear antibody escape from other sites. This approach was used to determine which sites to display in the logo plots, and which sites to include in the analysis of natural sequence variation.

### Data visualization

The static logo plot visualizations of the escape maps in the paper figures were created using the `dmslogo` package (https://jbloomlab.github.io/dmslogo/, version 0.3.2) and in all cases the height of each letter indicates the escape fraction for that amino-acid mutation calculated as described above. In Figure 2, we have separated the antibodies into two groups, and for each group the logo plots show all sites of escape from any antibody in that group according to the classification scheme described above. The code that generates these logo plot visualizations is available at https://github.com/jbloomlab/SARS-CoV-2-RBD_MAP_Crowe_antibodies/blob/master/results/summary/analyze_escape_profiles.md.

In many of the visualizations (*e*.*g*., Figure 2A), the RBD sites are categorized as falling into one of three structural regions (core RBD, RBM, or ACE2-contact residue) and colored accordingly. The RBM is defined as residues 437-508 (Li et al., 2005) with remaining residues comprising the core RBD. ACE2 contacts are defined as RBD residues with non-hydrogen atoms within 4 Angstrom of ACE2 atoms in the PDB: 6M0J crystal structure (Lan et al., 2020). In Figures 5B and S5, the letters in the escape maps are colored according to the effects of mutations on ACE2 binding or RBD expression as measured in Starr *et al*. (2020).

The multidimensional scaling in Figure 2D that projects the antibodies into a two-dimensional space of escape mutations was performed using the Python `scikit-learn` package. We first computed the similarity in the escape maps between each pair of antibodies as follows. Let *x*_*a*1_ be the vector of the total site escape values at each site for antibody *a1*. Then the similarity in escape between antibodies *a1* and *a2* is simply calculated as the dot product of the total site escape vectors after normalizing each vector to have a Euclidean norm of one; namely, the similarity is 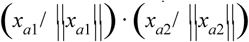. With this definition, the similarity is one if the total site escape is identical for the two antibodies, and zero if the escape is at completely distinct sites. We then calculated a dissimilarity for each pair of antibodies as simply one minus the similarity, and performed metric multidimensional scaling with two components on the dissimilarity matrix. The result is shown in Figure 2D, with antibodies shown in pie charts that are colored proportional to total squared site escape that falls into that RBD structural region. The code that generates these logo plot visualizations is available at https://github.com/jbloomlab/SARS-CoV-2-RBD_MAP_Crowe_antibodies/blob/master/results/summary/mds_escape_profiles.md.

For the static structural visualizations in the paper figures, the RBD surface (PDB: 6M0J, (Lan et al., 2020)) was colored by the largest-effect escape mutation at each site, with white indicating no escape and red indicating the strongest escape mutation for that antibody.

We created interactive structure-based visualizations of the escape maps using `dms-view` (Hilton et al., 2020) that are available at https://jbloomlab.github.io/SARS-CoV-2-RBD_MAP_Crowe_antibodies/. The logo plots in these escape maps can be colored according to the deep mutational scanning measurements of how mutations affect ACE2 binding or RBD expression as described above.

### Analysis of circulating variants and evolutionary conservation of antibody epitopes

All 94,233 spike sequences on GISAID as of 6 September 2020 were downloaded and aligned via `mafft` (Katoh and Standley, 2013). Sequences from non-human origins and sequences containing gap characters were removed, leaving 93,858 sequences. All RBD amino-acid mutations among GISAID sequences were enumerated, retaining only mutations that were sampled on at least one high-coverage sequence lacking undetermined ‘X’ characters within the RBD. All GISAID mutations at sites of escape from antibodies in our panel (using the method described above to define sites of escape) are shown in Figure S4. Counts were collapsed by site, and sites with at least 5 circulating mutations on GISAID are shown in Figure 5A. We acknowledge all GISAID contributors for their sharing of sequencing data (https://github.com/jbloomlab/SARS-CoV-2-RBD_MAP_Crowe_antibodies/blob/master/data/GISAID/gisaid_hcov-19_acknowledgement_table_2020_09_06.pdf).

To compute conservation of positions among sarbecoviruses, we used the RBD sequence set from Starr *et al*. (2020), which includes all unique RBD sequences curated by Letko *et al*. (2020), in addition to the non-Asian sarbecovirus BtKy72 (Tong et al., 2009) and newly described RBD sequences RaTG13 (Zhou et al., 2020b), RmYN02 (Zhou et al., 2020a), and GD-Pangolin and GX-Pangolin (Lam et al., 2020). RBD sequences were aligned at the amino-acid level via `mafft` with a gap opening penalty of 4.5. Alignment is available at https://github.com/jbloomlab/SARS-CoV-2-RBD_MAP_Crowe_antibodies/blob/master/data/RBDs_aligned.fasta. Shannon entropy of each alignment position was calculated using the `bio3d` package in R (Grant et al., 2006) as 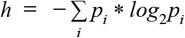, where *p*_*i*_ is the proportion of sequences with amino acid *i*. The effective number of amino acids at each position (*N*_eff_) was calculated as 2^*h*^.

### Pseudotyped lentiviral particles for neutralization assays and quantification of cellular entry

For neutralization assays, we used spike pseudotyped lentiviral particles that were generated essentially as described in Crawford *et al*. (2020a), using a codon-optimized SARS-CoV-2 spike from Wuhan-Hu-1 that contains a 21-amino-acid deletion at the end of the cytoplasmic tail that improves viral titers (Crawford et al., 2020b) along with the D614G mutation that is now prevalent in human SARS-CoV-2 (Korber et al., 2020). The plasmid encoding this spike, HDM_Spikedelta21_D614G, is available from Addgene (#158762), and the full sequence is at (https://www.addgene.org/158762/). Point mutations were introduced into the RBD of this plasmid via site-directed mutagenesis.

Pseudotyped lentiviral particles were generated as previously described (Crawford et al., 2020a). Viruses were rescued in biological duplicate (i.e., independent transfections). Briefly, 6e5 293T cells per well were seeded in 6-well plates in 2 mL D10 growth media (DMEM with 10% heat-inactivated FBS, 2 mM l-glutamine, 100 U/mL penicillin, and 100 μg/mL streptomycin). 24h later, cells were transfected using BioT transfection reagent (Bioland Scientific, Paramount, CA, USA) with a Luciferase_IRES_ZsGreen backbone, Gag/Pol lentiviral helper plasmid, and wildtype or mutant SARS-CoV-2 spike plasmids. Media was changed to fresh D10 at 24 h post-transfection. At 60 h post-transfection, viral supernatants were collected, filtered through a 0.45 μm SFCA low protein-binding filter, and stored at −80°C.

The resulting viruses were titered as previously described (Crawford et al., 2020a). 293T-ACE2 cells (BEI NR-52511) were seeded at 1.25e4 cells per well in 50 μL D10 in poly-L-lysine coated 96-well plates (Greiner 655930). After 24 h, 100 μL of diluted viral supernatants were added to cells across a dilution range of 4 serial 4-fold dilutions (*i*.*e*., 0.52 to 33.3 μL of virus were ultimately added to each well). Approximately 70 h post-infection, viral entry was quantified Bright-Glo Luciferase Assay System (Promega, E2610) as described in Crawford *et al*. (2020a). The relative titers reported in Figure S3D were calculated as the fold-change of relative luciferase units per microliter of each mutant RBD virus compared to unmutated RBD virus.

For the neutralization assays, the ACE2-293T cells were plated as described above for viral titering. 24 h later, pseudotyped lentivirus supernatants were diluted 1:6 and incubated with antibodies across a concentration range for 1 h at 37 °C, at a final concentration of antibody between 0.366 and 6,000 ng/mL. 100 μL of the virus-antibody mixture then was added to cells.

At ∼70 h post-infection, luciferase activity was measured as described above. Fraction infectivity of each antibody-containing well was calculated relative to a “no-antibody” well inoculated with the same initial viral supernatant (containing wildtype or mutant RBD) in the same row of the plate. We used the `neutcurve` package (https://jbloomlab.github.io/neutcurve/) to calculate the inhibitory concentration 50% (IC50) of each antibody against each virus by fitting a Hill curve with the bottom fixed at 0 and the top fixed at 1. The IC50 fold change relative to unmutated RBD was calculated for each mutant for each antibody.

### 293T mammalian cell-surface RBD display system

To validate the effects of individual mutations on antibody binding to the non-neutralizing antibody rCR3022 in a mammalian system as shown in Figure S3B,C, the RBD sequence used in yeast display was modified for mammalian surface display to create the HDM_Spike_RBD_B7-1 plasmid described in Loes *et al*. (2020). Site-directed mutagenesis was used to introduce single amino-acid substitutions into this plasmid.

293T cells were seeded at 6e5 cells per well in a 6-well plate. After 24 h, duplicate wells were transfected with 1 µg HDM_Spike_RBD_B7-1 plasmids and 1 µg of Transfection Carrier DNA (Promega, E4881) using BioT reagent (Bioland Sci, B01-02), according to manufacturer’s protocol. At 18 to 20 h post-transfection, cells were washed with phosphate buffered saline (PBS), dissociated from the plate with enzyme-free dissociation buffer (ThermoFisher, 13151014), harvested by centrifugation at 1,200 x *g* for 3 min, and washed in FACS buffer (PBS+1% bovine serum albumin). Cells were stained with recombinant biotinylated ACE2 (ACROBiosystems, AC2-H82E6) and serial dilutions of rCR3022 antibody for 1 h at room temperature, washed with FACS buffer, resuspended in a 1:200 dilution of PE-conjugated streptavidin (ThermoFisher, S866) and APC-conjugated Goat Anti-Human IgG (Jackson Labs, 109-115-098), and incubated on ice for 1 h. Cells were then washed twice in the FACS buffer and resuspended in PBS. rCR3022 antibody and ACE2-binding levels were determined via flow cytometry using a BD LSRFortessa X-50. 10,000 cells were analyzed at each rCR3022 concentration. Cells were gated to select for singleton events, ACE2 labeling was used to subset RBD+ cells and measure RBD expression, and rCR3022 labeling was measured within this RBD+ population. Compensation and gating was performed using FlowJo v10.7. EC50s were computed using the `neutcurve` package to fit four-parameter Hill curves (both baselines free) and the midpoint is reported as the EC50. The assays were performed on two separate days, and fold changes were computed relative to the unmutated (wildtype) RBD from that day.

### Production and purification of recombinant SARS-CoV-2 spike proteins for negative stain EM and binding competition experiments

We previously used a prefusion-stabilized, trimeric spike ectodomain (S2P_ecto_) to structurally define the sites several antibodies recognized on the SARS-CoV-2 spike trimer (Zost et al., 2020a). This construct is similar to ones previously reported (Wrapp et al., 2020) and includes the ectodomain of SARS-CoV-2 (to residue 1,208), a T4 fibritin trimerization domain, and C-terminal 8x-His tag and TwinStrep tags. The construct also includes K986P and V987P substitutions to stabilize the spike in the prefusion conformation and a mutated furin cleavage site. S2P_ecto_ protein was expressed in FreeStyle 293 cells (ThermoFisher) or Expi293 cells (ThermoFisher). Expressed S2P_ecto_ protein was isolated by metal affinity chromatography on HisTrap Excel columns (GE Healthcare), followed by further purification on a StrepTrap HP column (GE Healthcare) and size-exclusion chromatography on TSKgel G4000SW_XL_ (TOSOH). We also expressed a recently reported spike protein construct with 4 additional proline substitutions that enhance thermostability, yield, and structural homogeneity, here referred to as S6P_ecto_ (Hsieh et al., 2020). The S6P_ecto_ protein was expressed in FreeStyle293 cells and isolated on a StrepTrap HP column following the addition of BioLock Biotin Blocking Solution (IBA Lifesciences) to the culture supernatant.

### Negative stain electron microscopy of SARS-CoV-2 S/Fab complexes

Fabs were produced for negative stain electron microscopy by digesting recombinant chromatography-purified IgGs using resin-immobilized cysteine protease enzyme (FabALACTICA, Genovis). Digestions were performed in 100 mM sodium phosphate, 150 mM NaCl pH 7.2 (PBS) for ∼16 h at ambient temperature. After digestion, the digestion mix was incubated with CaptureSelect Fc resin (Genovis) for 30 min at ambient temperature in PBS buffer to remove cleaved Fc and intact, undigested IgG. If needed, the Fab was buffer exchanged into Tris buffer by centrifugation with a Zeba spin column (Thermo Scientific).

For screening and imaging of negatively-stained (NS) SARS-CoV-2 S2P_ecto_ or SARS-CoV-2 S6P_ecto_ protein in complex with human Fabs, the proteins were incubated at a molar ratio of 4 Fab:3 spike monomer for ∼1 h and approximately 3 µL of the sample at concentrations of about 10 to 15 µg/mL was applied to a glow discharged grid with continuous carbon film on 400 square mesh copper EM grids (Electron Microscopy Sciences). The grids were stained with 0.75% uranyl formate (UF) (Ohi et al., 2004). Images were collected using a Gatan US4000 4k × 4k CCD camera on a FEI TF20 (TFS) transmission electron microscope operated at 200 keV and controlled with SerialEM (Mastronarde, 2005). All images were taken at 50,000x magnification with a pixel size of 2.18 Å/pix in low-dose mode at a defocus of 1.5 to 1.8 μm.

The total dose for the micrographs was ∼25 to 38 e^−^/Å^2^. Image processing was performed using the cryoSPARC software package (Punjani et al., 2017). Images were imported, and the micrographs were CTF estimated. The images then were picked with Topaz (Bepler et al., 2019, 2020). The particles were extracted with a box size of 256 pixels and binned to 128 pixels giving pixel size of 4.36 Å/pix. 2D class averages were performed and good classes selected for *ab-initio* model and refinement without symmetry. For EM model docking of SARS-CoV-2 S complexed with Fabs, the “RBD up” structure of SARS-CoV-2 (PDB: 6VYB) (Walls et al., 2020b) and a Fab crystal structure (Fab: 12E8) were used in Chimera (see Table S1 for details). To visualize escape maps on the SARS-CoV-2 trimer, the crystal structure of SARS-CoV-2 RBD (solved in complex with ACE2, PDB: 6M0J) was aligned to the RBD in the cryo EM density of trimeric spike. All images were made with Chimera.

### Antibody competition-binding analysis

For the competition experiments reported in Figure S6D, wells of 384-well microtiter plates were coated with 1 µg/mL of purified SARS-CoV-2 S6P_ecto_ protein at 4°C overnight. Plates were blocked with 2% BSA in DPBS containing 0.05% Tween-20 (DPBS-T) for 1 h. Purified unlabeled antibodies were diluted to 20 µg/mL in blocking buffer, added to the wells (20 μL/well) in triplicate, and incubated for 1 h at ambient temperature. SARS-CoV-2 antibodies were added to each of three wells with the respective antibody at 2.5 μg/mL in a 5 μL/well volume (final 0.1 μg/mL concentration of biotinylated antibody) without washing of unlabeled antibody and then incubated for 1 h at ambient temperature. Plates were washed, and bound antibodies were detected using HRP-conjugated avidin (Sigma) and TMB substrate. The signal obtained for binding of the biotin-labeled reference antibody in the presence of the unlabeled tested antibody was expressed as a percentage of the binding of the reference antibody alone after subtracting the background signal. Tested antibodies were considered competing if their presence reduced the reference antibody binding to less than 30 % of its maximal binding and non-competing if the signal was greater than 75%. A level of 30–75% was considered intermediate competition.

### VSV viruses expressing SARS-CoV-2 spike protein

The generation of a replication-competent vesicular stomatitis virus (VSV) expressing SARS-CoV-2 S protein that replaces VSV G protein (VSV-SARS-CoV-2) has been described previously (Case et al., 2020). This virus encodes the spike protein from SARS-CoV-2 with a 21 amino-acid C-terminal deletion. The spike-expressing VSV virus was propagated in MA104 cells (ATCC CRL-2378.1) as described previously (Case et al., 2020), and viral stocks were titrated on Vero E6 cell monolayer cultures. Plaques were visualized using crystal violet staining.

### Selection of escape mutants using the spike-expressing VSV

To screen for escape mutations selected in the presence of individual antibodies or antibody cocktails, we used a real-time cell analysis assay (RTCA) and xCELLigence RTCA MP Analyzer (ACEA Biosciences Inc.) with modification of previously described assays (Gilchuk et al., 2020a; Weisblum et al., 2020). Fifty (50) μL of cell culture medium (DMEM supplemented with 2% FBS) was added to each well of a 96-well E-plate to obtain a background reading. Eighteen thousand (18,000) Vero E6 cells in 50 μL of cell culture medium were seeded per each well, and plates were placed on the analyzer. Measurements were taken automatically every 15 min and the sensograms were visualized using RTCA software version 2.1.0 (ACEA Biosciences Inc). VSV-SARS-CoV-2 virus (500 plaque forming units [PFU] per well, ∼0.03 MOI) was mixed with a saturating neutralizing concentration of individual antibody (5 µg/mL) or two-antibody cocktail (1:1 antibody ratio, 5 μg/mL total antibody concentration) in a total volume of 100 μL and incubated for 1 h at 37°C. At 16-20 h after seeding the cells, the virus-antibody mixtures were added into 8 to 96 replicate wells of 96-well E-plates with cell monolayers. Wells containing only virus in the absence of antibody and wells containing only Vero E6 cells in medium were included on each plate as controls. Plates were measured continuously (every 15 min) for 72 h. The escape mutants were identified by delayed CPE in wells containing antibody. To verify escape from antibody selection, isolated viruses were assessed in a subsequent RTCA experiment in the presence of 10 μg/mL of mAb as used for the escape virus selection and a partner mAb recognizing non-overlapping epitope residues (see Figure 6A).

### Sequence analysis of the gene encoding spike protein from spike protein-expressing VSV escape mutants

To identify escape mutations present in spike protein-expressing VSV antibody-selected escape variants, the escape viruses isolated after RTCA escape screening were propagated in 6-well culture plates with confluent Vero E6 cells in the presence of 10 μg/mL of the corresponding antibody. Viral RNA was isolated using a QiAmp Viral RNA extraction kit (Qiagen) from aliquots of supernatant containing a suspension of the selected virus population. The spike protein gene cDNA was amplified with a SuperScript IV One-Step RT-PCR kit (ThermoFisher Scientific) using primers flanking the S gene. The amplified PCR product (∼ 4,000 bp) was purified using SPRI magnetic beads (Beckman Coulter) at a 1:1 ratio and sequenced by the Sanger sequence technique using primers giving forward and reverse reads of the RBD.

## Supplemental Figures

**Figure S1.**
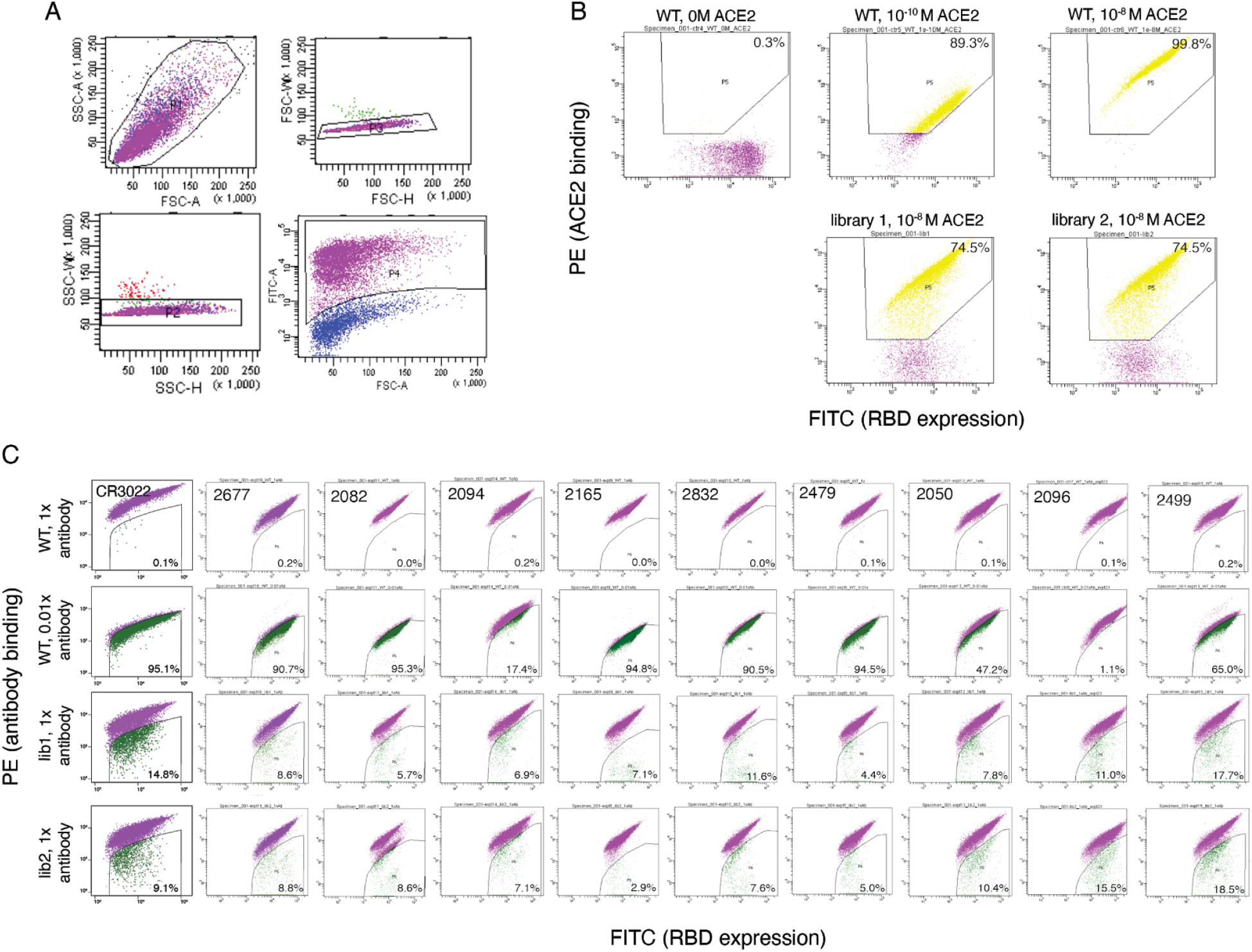
FACS gating. (A) Representative hierarchical gates drawn to isolate RBD+ single cells as the parent population for FACS gates in (B, C). First, hierarchical gates were drawn to select single-cell events: forward scatter (FSC) versus side scatter (SSC, top left), SSC width versus height (bottom left), and FSC width versus height (top right). Next, FITC+ labeling of a C-terminal epitope tag on the RBD was used to identify RBD+ cells (purple, bottom right). Selection gates for ACE2+ and antibody-negative sorts (B, C) are nested within this RBD+ population. (B) RBD mutant libraries were first sorted for variants that could bind ACE2 with at least 0.01x the affinity of unmutated SARS-CoV-2 RBD. Top three plots show unmutated SARS-CoV-2 labeled at 0 M, 1e-10 M, and 1e-8 M ACE2. A selection gate was drawn to capture unmutated cells labeled at 1e-10 M ACE2. The bottom two plots show the application of this selection gate to the duplicate RBD mutant libraries labeled at 1e-8 M ACE2. Percentages of RBD+ cells (yellow) in each control and library sample that fall into the ACE2+ sort bin are shown in the upper-right of each FACS plot. These ACE2+ sorted libraries were grown overnight and used for subsequent antibody-escape selections. (C) Selection gates for the antibody-escape sorts. Unmutated SARS-CoV-2 RBD was labeled at 400 ng/mL (1x) and 4 ng/mL (0.01x) with each antibody. Antibody-escape selection gates were drawn to capture 0.2% or less of the 1x and up to 95% of the 0.01x antibody-labeled unmutated RBD control cells. Each mutant RBD library was labeled with 400 ng/mL (1x) antibody, and cells that were captured in the “antibody-escape bin” were sorted and their barcodes were sequenced. Percentages of RBD+ cells in each control and library sample that fall into the antibody-escape bin are shown in the bottom-right of each FACS plot.

**Figure S2.**
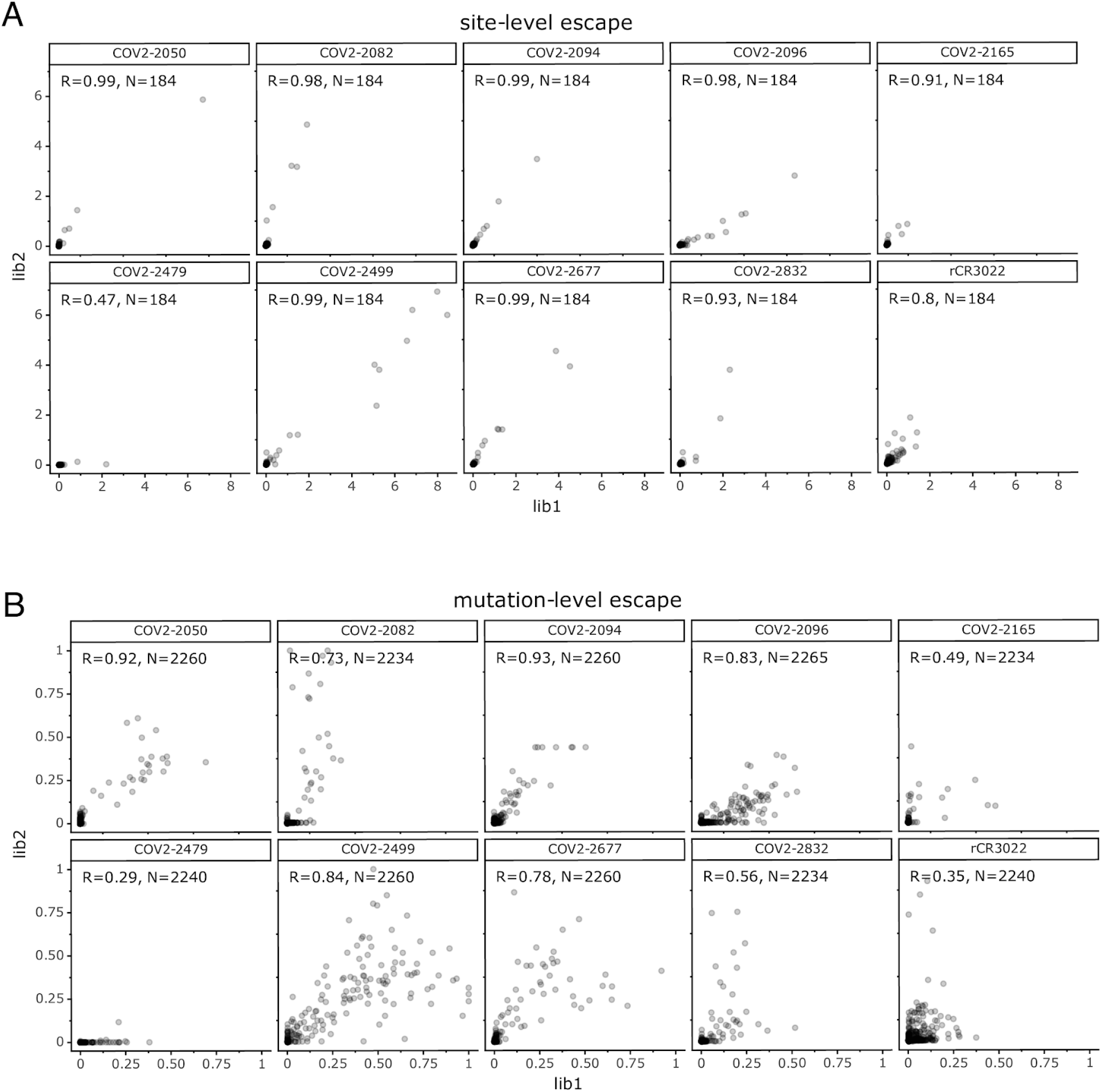
Correlation between the duplicate mappings of escape mutations made with the independently generated mutant virus libraries (“lib1” and “lib2”). (A) Correlation between the total escape at each site. (B) Correlation between the escape fraction measured for each individual mutation. The text insets in each plot give the Pearson’s correlation coefficient and the number of sites or mutations for which measurements were made for both libraries. The data shown in the rest of the paper are the average of those from the two libraries.

**Figure S3.**
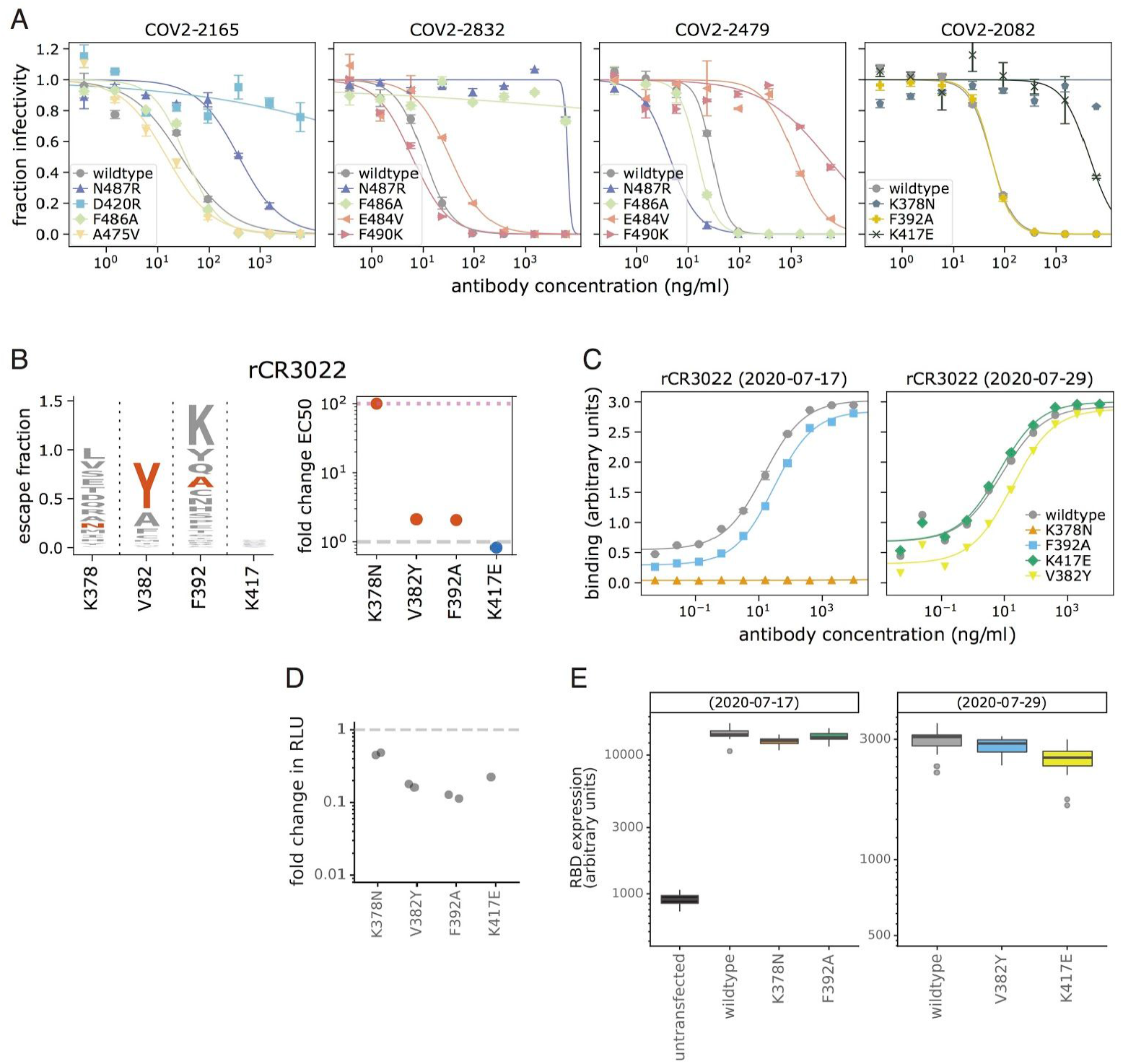
Full curves for validation neutralization assays, and effects of mutations on antibody binding to mammalian-expressed rCR3022. (A) Neutralization curves with the spike-pseudotyped lentiviral particles used to determine IC50 values plotted in Figure 3. Each point represents the mean and standard error of 2 independent measurements. The IC50s were computed using the `neutcurve` package (https://jbloomlab.github.io/neutcurve/) to fit two-parameter Hill curves (with the baselines fixed to 0 and 1). IC50s outside the range of tested antibody concentrations are reported as upper bounds. (B) Antibody rCR3022 is non-neutralizing, so we instead used flow cytometry to measure rCR3022 binding to RBD expressed on the surface of mammalian cells (see Methods for details), with the values representing the fold change in effective concentration 50% (EC50) for antibody binding to each mutant RBD relative to wildtype. (C) The binding curves summarized in (B), with the y-axis representing binding as measured by flow cytometry. EC50s are computed using the `neutcurve` package to fit four-parameter Hill curves (both baselines free) and the midpoint is reported as the EC50. The assays were performed on two separate days, and fold changes are computed relative to the unmutated (wildtype) RBD from that day. (D) rCR3022 escape mutations are compatible with function in spike-pseudotyped lentiviral particles. The infectious titer of spike-pseudotyped lentivirus mutants in transfection supernatants as quantified by fold change in relative luciferase units (RLUs) compared to virus pseudotyped with the unmutated (wildtype) spike. All titers were measured in biological duplicate transfections (two jittered points) except K417E. (E) To estimate RBD expression on the surface of 293T cells in the rCR3022 binding assays in panels B and C, cells were also labeled with biotinylated ACE2 and fluorophore-conjugated streptavidin. ACE2 binding levels, a proxy for RBD expression, were measured by flow cytometry. Box plots represent the median and 25th and 75th percentiles, whiskers are 1.5 * interquartile range, and outliers are shown individually. For each condition, n=12-24.

**Figure S4.**
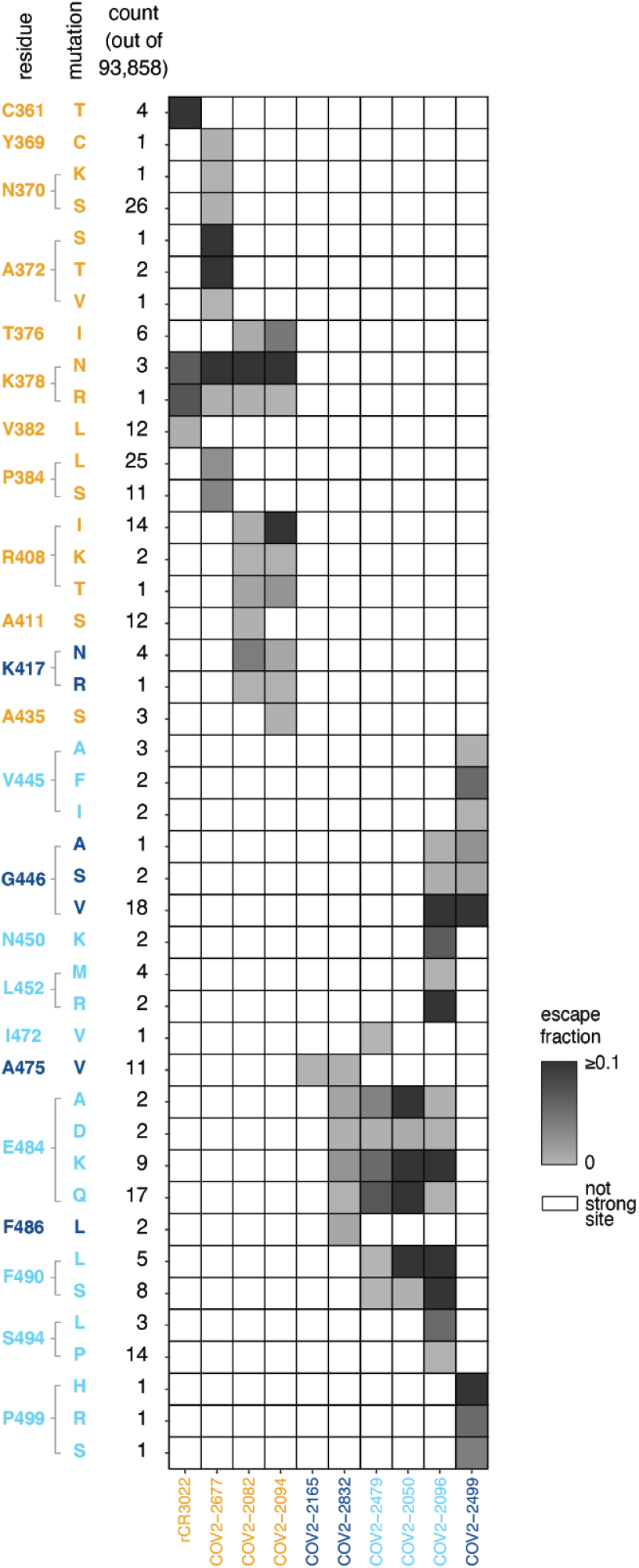
Variation at sites of antibody escape among currently circulating SARS-CoV-2 viruses. Table shows all RBD mutations sampled among sequences in GISAID as of 6 September 2020 at sites of escape from at least one antibody. Cells are colored by escape fraction of the individual circulating mutant for each antibody: white cells indicate sites that are not sites of escape from an antibody; for sites of escape, per-mutation escape fraction is colored from light to dark gray, with any mutation conferring >0.1 escape fraction colored equally dark. Sites are in orange for the core RBD, light blue for the RBM, and dark blue for ACE2 contact residues. Antibodies are colored according to where the majority of their sites of escape fall. These per-mutation counts are collapsed into the site-wise table presented in Figure 5A.

**Figure S5.**
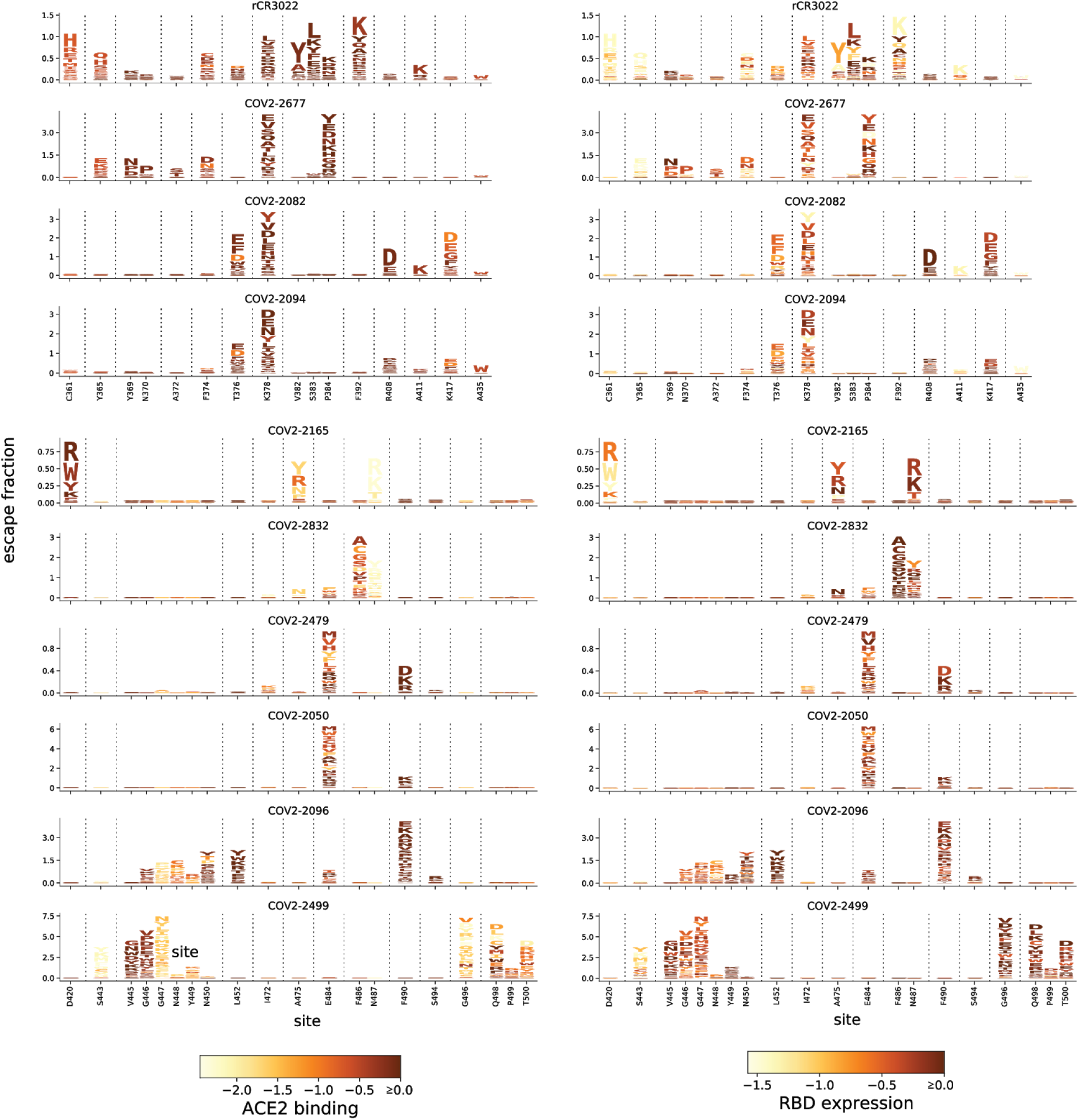
Logo plots of antibody escape accounting for mutation effects on ACE2-binding affinity and RBD folding. Logo plots as in Figure 2C. Mutations are colored according to their effects on ACE2-binding affinity (left) or RBD folding and expression (right), as measured previously (Starr et al., 2020). Some mutations annotated as escape in our main display impair ACE2 binding or RBD folding, which may limit their fitness in the context of virus particles.

**Figure S6.**
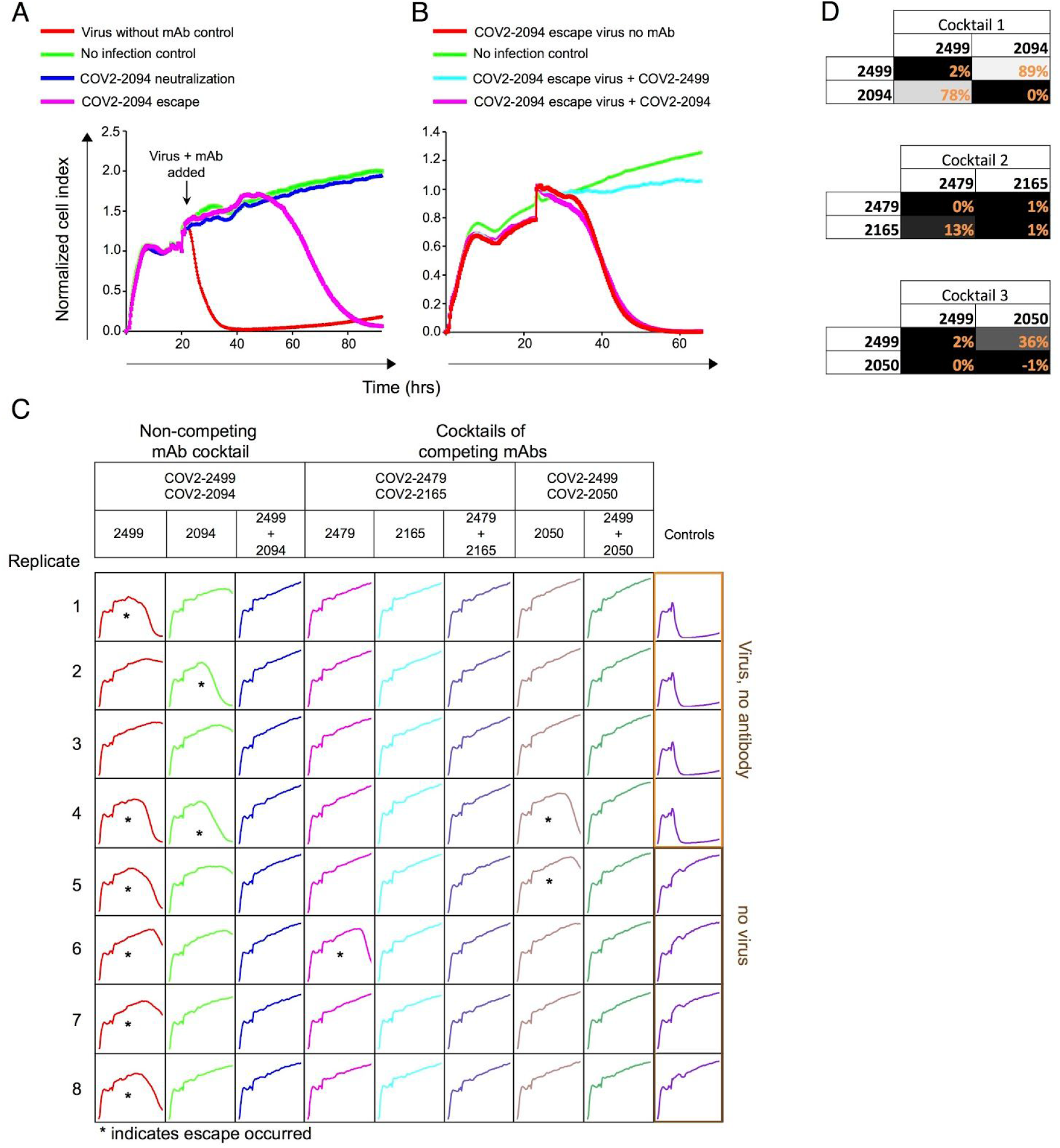
Real-time cell analysis (RTCA) to select for spike-expressing VSV viruses that escape antibody neutralization, and antibody competition for binding to RBD. (A) Representative RTCA sensograms showing virus that escaped antibody neutralization. Cytopathic effect (CPE) was monitored kinetically in Vero E6 cells inoculated with virus in the presence of a saturating concentration of antibody COV2-2094 (5 µg/mL). Escape (magenta) or lack of escape (blue) are shown. Uninfected cells (green) or cells inoculated with virus without antibody (red) serve as controls. Magenta and blue curves represent a single representative well; the red and green controls are mean of technical quadruplicates. (B) Representative RTCA sensograms validating that the virus selected by COV2-2094 in panel (A) indeed escaped COV2-2094 (magenta) but was neutralized by COV2-2499 (light blue). (C) Example sensograms from individual wells of 96-well E-plate analysis showing viruses that escaped neutralization (noted with *) by indicated antibodies. (D) Competition assays for RBD binding, with percentages showing binding of a second labeled antibody to the RBD after pre-binding with the first antibody. Values close to 0% indicate complete competition, and values close to 100% indicate lack of competition.

**Table S1.** Summary of electron microscopy data collection and statistics for SARS-CoV-2 S protein in complex with human Fabs

|  |  |  |  |  |  |  |
| --- | --- | --- | --- | --- | --- | --- |
|  |  | Structure of SARS-CoV-2 S2P<sub>ecto</sub> or S6P<sub>ecto</sub> proteins in complex with indicated Fabs |  |  |  |  |
|  |  | Fab COV2-2082 | Fab COV2-2096 | Fab COV2-2165<br>* | Fab COV2-2479 | Fab COV2-2832 |
|  | EMDB #: | EMD-22627 | EMD-22148 | EMD-21974 | EMD-22628 | EMD-22149 |
| Microscope setting | Microscope | TF-20 | TF-20 | TF-20 | TF-20 | TF-20 |
|  | Voltage (kV) | 200 | 200 | 200 | 200 | 200 |
|  | Detector | US-4000 CCD | US-4000 CCD | US-4000 CCD | US-4000 CCD | US-4000 CCD |
| | Magnification | 50,000× | 50,000× | 50,000× | 50,000× | 50,000× |
|  | Pixel size | 2.18 | 2.18 | 2.18 | 2.18 | 2.18 |
|  | Exposure (e-/Å<sup>2</sup>) | 30 | 30 | 25 | 30 | 30 |
|  | Defocus range (μm) | 1.5 to 1.8 | 1.5 to 1.8 | 1.5 to 1.8 | 1.5 to 1.8 | 1.5 to 1.8 |
| Data | Antigen | S6P<sub>ecto</sub> | S2P<sub>ecto</sub> | S2P<sub>ecto</sub> | S6P<sub>ecto</sub> | S2P<sub>ecto</sub> |
|  | Micrographs, # | 237 | 562 | 83 | 331 | 514 |
|  | Particles, # | 972 | 19,728 | 3,705 | 81,758 | 7,773 |
|  | Particles #, after 2D | 673 | 18,202 | 1,868 | 76,431 | 3,778 |
|  | Final particles, # | 663 | 12,132 | 1,057 | 18,535 | 3,424 |
|  | Symmetry | C1 | C1 | C1 | C1 | C1 |
| Model docking | CoV-2-S CC | PDB: 6VYB 0.895 | PDB: 6VYB 0.836 | PDB: 6VYB 0.828 | PDB: 6VYB 0.900 | PDB: 6VYB 0.8952 |
|  | Fab (PDB: 12E8) CC | 0.91 | 0.916 | 0.905 | 0.91 | 0.913 |

\*Previously reported (Zost *et al.*, 2020a)

**Supplemental File S1:** The estimates of the antigenic effects of all mutations for all antibodies. The file gives the “escape fraction” for each mutation, as well as the total escape fraction at each site and the maximum escape fraction for any mutation at the site. The file is also available on GitHub at https://raw.githubusercontent.com/jbloomlab/SARS-CoV-2-RBD_MAP_Crowe_antibodies/master/results/supp_data/MAP_paper_antibodies_raw_data.csv.

## Notes

https://jbloomlab.github.io/SARS-CoV-2-RBD_MAP_Crowe_antibodies/

### Summary of Updates

Updated acknowledgements to include additional funding sources.

## References

Addetia, A., Crawford, K.H.D., Dingens, A., Zhu, H., Roychoudhury, P., Huang, M.-L., Jerome, K.R., Bloom, J.D., and Greninger, A.L. (2020). Neutralizing antibodies correlate with protection from SARS-CoV-2 in humans during a fishery vessel outbreak with high attack rate. J. Clin. Microbiol.

Alsoussi, W.B., Turner, J.S., Case, J.B., Zhao, H., Schmitz, A.J., Zhou, J.Q., Chen, R.E., Lei, T., Rizk, A.A., McIntire, K.M., et al. (2020). A potently neutralizing antibody protects mice against SARS-CoV-2 infection. J. Immunol. 205, 915–922.

Amir, E.-A.D., Davis, K.L., Tadmor, M.D., Simonds, E.F., Levine, J.H., Bendall, S.C., Shenfeld, D.K., Krishnaswamy, S., Nolan, G.P., and Pe’er, D. (2013). viSNE enables visualization of high dimensional single-cell data and reveals phenotypic heterogeneity of leukemia. Nat. Biotechnol. 31, 545–552.

Barnes, C.O., West, A.P., Jr, Huey-Tubman, K.E., Hoffmann, M.A.G., Sharaf, N.G., Hoffman, P.R., Koranda, N., Gristick, H.B., Gaebler, C., Muecksch, F., et al. (2020a). Structures of human antibodies bound to SARS-CoV-2 Spike reveal common epitopes and recurrent features of antibodies. Cell 182, 828–842.e16.

Barnes, C.O., Jette, C.A., Abernathy, M.E., Dam, K.-M.A., Esswein, S.R., Gristick, H.B., Malyutin, A.G., Sharaf, N.G., Huey-Tubman, K.E., Lee, Y.E., et al. (2020b). Structural classification of neutralizing antibodies against the SARS-CoV-2 spike receptor-binding domain suggests vaccine and therapeutic strategies. bioRxiv 2020.08.30.273920.

Baum, A., Fulton, B.O., Wloga, E., Copin, R., Pascal, K.E., Russo, V., Giordano, S., Lanza, K., Negron, N., Ni, M., et al. (2020a). Antibody cocktail to SARS-CoV-2 spike protein prevents rapid mutational escape seen with individual antibodies. Science 369, 1014–1018.

Baum, A., Copin, R., Ajithdoss, D., Zhou, A., Lanza, K., Negron, N., Ni, M., Wei, Y., Atwal, G.S., Oyejide, A., et al. (2020b). REGN-COV2 antibody cocktail prevents and treats SARS-CoV-2 infection in rhesus macaques and hamsters. bioRxiv.

Becht, E., McInnes, L., Healy, J., Dutertre, C.-A., Kwok, I.W.H., Ng, L.G., Ginhoux, F., and Newell, E.W. (2019). Dimensionality reduction for visualizing single-cell data using UMAP. Nat. Biotechnol. 37.

Bepler, T., Morin, A., Rapp, M., Brasch, J., Shapiro, L., Noble, A.J., and Berger, B. (2019). Positive-unlabeled convolutional neural networks for particle picking in cryo-electron micrographs. Nat. Methods 16, 1153–1160.

Bepler, T., Kelley, K., Noble, A.J., and Berger, B. (2020). Topaz-Denoise: general deep denoising models for cryoEM and cryoET. bioRxiv.

Bogan, A.A., and Thorn, K.S. (1998). Anatomy of hot spots in protein interfaces. J. Mol. Biol. 280, 1–9.

Brouwer, P.J.M., Caniels, T.G., van der Straten, K., Snitselaar, J.L., Aldon, Y., Bangaru, S., Torres, J.L., Okba, N.M.A., Claireaux, M., Kerster, G., et al. (2020). Potent neutralizing antibodies from COVID-19 patients define multiple targets of vulnerability. Science 369, 643–650.

Cao, Y., Su, B., Guo, X., Sun, W., Deng, Y., Bao, L., Zhu, Q., Zhang, X., Zheng, Y., Geng, C., et al. (2020). Potent neutralizing antibodies against SARS-CoV-2 identified by high-throughput single-cell sequencing of convalescent patients’ B cells. Cell 182, 73–84.e16.

Case, J.B., Rothlauf, P.W., Chen, R.E., Liu, Z., Zhao, H., Kim, A.S., Bloyet, L.-M., Zeng, Q., Tahan, S., Droit, L., et al. (2020). Neutralizing antibody and soluble ACE2 inhibition of a replication-competent VSV-SARS-CoV-2 and a clinical isolate of SARS-CoV-2. Cell Host Microbe.

Chen, W.-H., Du, L., Chag, S.M., Ma, C., Tricoche, N., Tao, X., Seid, C.A., Hudspeth, E.M., Lustigman, S., Tseng, C.-T.K., et al. (2014). Yeast-expressed recombinant protein of the receptor-binding domain in SARS-CoV spike protein with deglycosylated forms as a SARS vaccine candidate. Hum. Vaccin. Immunother. 10, 648–658.

Clackson, T., and Wells, J.A. (1995). A hot spot of binding energy in a hormone-receptor interface. Science 267, 383–386.

Crawford, K.H.D., Eguia, R., Dingens, A.S., Loes, A.N., Malone, K.D., Wolf, C.R., Chu, H.Y., Tortorici, M.A., Veesler, D., Murphy, M., et al. (2020a). Protocol and reagents for pseudotyping lentiviral particles with SARS-CoV-2 Spike protein for neutralization assays. Viruses 12.

Crawford, K.H.D., Dingens, A.S., Eguia, R., Wolf, C.R., Wilcox, N., Logue, J.K., Shuey, K., Casto, A.M., Fiala, B., Wrenn, S., et al. (2020b). Dynamics of neutralizing antibody titers in the months after SARS-CoV-2 infection (medRxiv).

Cunningham, B.C., and Wells, J.A. (1993). Comparison of a structural and a functional epitope. J. Mol. Biol. 234, 554–563.

Dall’Acqua, W., Goldman, E.R., Lin, W., Teng, C., Tsuchiya, D., Li, H., Ysern, X., Braden, B.C., Li, Y., Smith-Gill, S.J., et al. (1998). A mutational analysis of binding interactions in an antigen-antibody protein-protein complex. Biochemistry 37, 7981–7991.

Dieterle, M.E., Haslwanter, D., Bortz, R.H., 3rd, Wirchnianski, A.S., Lasso, G., Vergnolle, O., Abbasi, S.A., Fels, J.M., Laudermilch, E., Florez, C., et al. (2020). A replication-competent vesicular stomatitis virus for studies of SARS-CoV-2 Spike-mediated cell entry and its inhibition. Cell Host Microbe.

Dingens, A.S., Arenz, D., Weight, H., Overbaugh, J., and Bloom, J.D. (2019). An antigenic atlas of HIV-1 escape from broadly neutralizing antibodies distinguishes functional and structural epitopes. Immunity 50, 520–532.e3.

Gilchuk, P., Murin, C.D., Milligan, J.C., Cross, R.W., Mire, C.E., Ilinykh, P.A., Huang, K., Kuzmina, N., Altman, P.X., Hui, S., et al. (2020a). Analysis of a therapeutic antibody cocktail reveals determinants for cooperative and broad ebolavirus neutralization. Immunity 52, 388–403.e12.

Gilchuk, P., Bombardi, R.G., Erasmus, J.H., Tan, Q., Nargi, R., Soto, C., Abbink, P., Suscovich, T.J., Durnell, L.A., Khandhar, A., et al. (2020b). Integrated pipeline for the accelerated discovery of antiviral antibody therapeutics. Nat Biomed Eng.

Grant, B.J., Rodrigues, A.P.C., ElSawy, K.M., McCammon, J.A., and Caves, L.S.D. (2006). Bio3d: an R package for the comparative analysis of protein structures. Bioinformatics 22, 2695–2696.

Hamilton, S.R., Bobrowicz, P., Bobrowicz, B., Davidson, R.C., Li, H., Mitchell, T., Nett, J.H., Rausch, S., Stadheim, T.A., Wischnewski, H., et al. (2003). Production of complex human glycoproteins in yeast. Science 301, 1244–1246.

Hassan, A.O., Case, J.B., Winkler, E.S., Thackray, L.B., Kafai, N.M., Bailey, A.L., McCune, B.T., Fox, J.M., Chen, R.E., Alsoussi, W.B., et al. (2020). A SARS-CoV-2 infection model in mice demonstrates protection by neutralizing antibodies. Cell 182, 744–753.e4.

He, B., Zhang, Y., Xu, L., Yang, W., Yang, F., Feng, Y., Xia, L., Zhou, J., Zhen, W., Feng, Y., et al. (2014). Identification of diverse alphacoronaviruses and genomic characterization of a novel severe acute respiratory syndrome-like coronavirus from bats in China. J. Virol. 88, 7070–7082.

Hilton, S., Huddleston$, J., Black, A., North, K., Dingens, A., Bedford, T., and Bloom, J. (2020). dms-view: Interactive visualization tool for deep mutational scanning data. JOSS 5, 2353.

Hsieh, C.-L., Goldsmith, J.A., Schaub, J.M., DiVenere, A.M., Kuo, H.-C., Javanmardi, K., Le, K.C., Wrapp, D., Lee, A.G., Liu, Y., et al. (2020). Structure-based design of prefusion-stabilized SARS-CoV-2 spikes. Science.

Huang, K.-Y.A., Tan, T., Chen, T.-H., Huang, C.-G., Harvey, R., Hussain, S., Chen, C.-P., Harding, A., Gilbert-Jaramillo, J., Liu, X., et al. (2020). Plasmablast-derived antibody response to acute SARS-CoV-2 infection in humans.

Huo, J., Zhao, Y., Ren, J., Zhou, D., Duyvesteyn, H.M.E., Ginn, H.M., Carrique, L., Malinauskas, T., Ruza, R.R., Shah, P.N.M., et al. (2020). Neutralization of SARS-CoV-2 by destruction of the prefusion Spike. Cell Host Microbe.

Jin, L., Fendly, B.M., and Wells, J.A. (1992). High resolution functional analysis of antibody-antigen interactions. J. Mol. Biol. 226, 851–865.

Ju, B., Zhang, Q., Ge, J., Wang, R., Sun, J., Ge, X., Yu, J., Shan, S., Zhou, B., Song, S., et al. (2020). Human neutralizing antibodies elicited by SARS-CoV-2 infection. Nature 584, 115–119.

Julg, B., Liu, P.-T., Wagh, K., Fischer, W.M., Abbink, P., Mercado, N.B., Whitney, J.B., Nkolola, J.P., McMahan, K., Tartaglia, L.J., et al. (2017). Protection against a mixed SHIV challenge by a broadly neutralizing antibody cocktail. Sci. Transl. Med. 9.

Katoh, K., and Standley, D.M. (2013). MAFFT multiple sequence alignment software version 7: improvements in performance and usability. Mol. Biol. Evol. 30, 772–780.

Korber, B., Fischer, W.M., Gnanakaran, S., Yoon, H., Theiler, J., Abfalterer, W., Hengartner, N., Giorgi, E.E., Bhattacharya, T., Foley, B., et al. (2020). Tracking changes in SARS-CoV-2 Spike: evidence that D614G increases infectivity of the COVID-19 virus. Cell 182, 812–827.e19.

Lam, T.T.-Y., Jia, N., Zhang, Y.-W., Shum, M.H.-H., Jiang, J.-F., Zhu, H.-C., Tong, Y.-G., Shi, Y.-X., Ni, X.-B., Liao, Y.-S., et al. (2020). Identifying SARS-CoV-2-related coronaviruses in Malayan pangolins. Nature 583, 282–285.

Lan, J., Ge, J., Yu, J., Shan, S., Zhou, H., Fan, S., Zhang, Q., Shi, X., Wang, Q., Zhang, L., et al. (2020). Structure of the SARS-CoV-2 spike receptor-binding domain bound to the ACE2 receptor. Nature 581, 215–220.

Lee, J.M., Eguia, R., Zost, S.J., Choudhary, S., Wilson, P.C., Bedford, T., Stevens-Ayers, T., Boeckh, M., Hurt, A.C., Lakdawala, S.S., et al. (2019). Mapping person-to-person variation in viral mutations that escape polyclonal serum targeting influenza hemagglutinin. Elife 8.

Letko, M., Marzi, A., and Munster, V. (2020). Functional assessment of cell entry and receptor usage for SARS-CoV-2 and other lineage B betacoronaviruses. Nat Microbiol 5, 562–569.

Li, F., Li, W., Farzan, M., and Harrison, S.C. (2005). Structure of SARS coronavirus spike receptor-binding domain complexed with receptor. Science 309, 1864–1868.

Li, Q., Wu, J., Nie, J., Zhang, L., Hao, H., Liu, S., Zhao, C., Zhang, Q., Liu, H., Nie, L., et al. (2020). The impact of mutations in SARS-CoV-2 Spike on viral infectivity and antigenicity. Cell 182, 1284–1294.e9.

Linnemann, C.C., Jr (1973). Measles vaccine: immunity, reinfection and revaccination. Am. J. Epidemiol. 97, 365–371.

Liu, L., Wang, P., Nair, M.S., Yu, J., Rapp, M., Wang, Q., Luo, Y., Chan, J.F.-W., Sahi, V., Figueroa, A., et al. (2020). Potent neutralizing antibodies against multiple epitopes on SARS-CoV-2 spike. Nature 584, 450–456.

Loes, A.N., Gentles, L.E., Greaney, A.J., Crawford, K.H.D., and Bloom, J.D. (2020). Attenuated influenza virions expressing the SARS-CoV-2 receptor-binding domain induce neutralizing antibodies in mice. Viruses 12, 987.

Mastronarde, D.N. (2005). Automated electron microscope tomography using robust prediction of specimen movements. J. Struct. Biol. 152, 36–51.

ter Meulen, J., van den Brink, E.N., Poon, L.L.M., Marissen, W.E., Leung, C.S.W., Cox, F., Cheung, C.Y., Bakker, A.Q., Bogaards, J.A., van Deventer, E., et al. (2006). Human monoclonal antibody combination against SARS coronavirus: synergy and coverage of escape mutants. PLoS Med. 3, e237.

Ohi, M., Li, Y., Cheng, Y., and Walz, T. (2004). Negative staining and image classification - powerful tools in modern electron microscopy. Biol. Proced. Online 6, 23–34.

Otwinowski, J., McCandlish, D.M., and Plotkin, J.B. (2018). Inferring the shape of global epistasis. Proc. Natl. Acad. Sci. U. S. A. 115, E7550–E7558.

Panum, P.L. (1939). Observation made during the epidemic of measles on the Faroe Islands in the year 1846. Med Classics 3, 839–886.

Punjani, A., Rubinstein, J.L., Fleet, D.J., and Brubaker, M.A. (2017). cryoSPARC: algorithms for rapid unsupervised cryo-EM structure determination. Nat. Methods 14, 290–296.

Robbiani, D.F., Gaebler, C., Muecksch, F., Lorenzi, J.C.C., Wang, Z., Cho, A., Agudelo, M., Barnes, C.O., Gazumyan, A., Finkin, S., et al. (2020). Convergent antibody responses to SARS-CoV-2 in convalescent individuals. Nature 584, 437–442.

Rockx, B., Donaldson, E., Frieman, M., Sheahan, T., Corti, D., Lanzavecchia, A., and Baric, R.S. (2010). Escape from human monoclonal antibody neutralization affects in vitro and in vivo fitness of severe acute respiratory syndrome coronavirus. J. Infect. Dis. 201, 946–955.

Rogers, T.F., Zhao, F., Huang, D., Beutler, N., Burns, A., He, W.-T., Limbo, O., Smith, C., Song, G., Woehl, J., et al. (2020). Isolation of potent SARS-CoV-2 neutralizing antibodies and protection from disease in a small animal model. Science 369, 956–963.

Schmidt, A.G., Therkelsen, M.D., Stewart, S., Kepler, T.B., Liao, H.-X., Moody, M.A., Haynes, B.F., and Harrison, S.C. (2015). Viral receptor-binding site antibodies with diverse germline origins. Cell 161, 1026–1034.

Schommers, P., Gruell, H., Abernathy, M.E., Tran, M.-K., Dingens, A.S., Gristick, H.B., Barnes, C.O., Schoofs, T., Schlotz, M., Vanshylla, K., et al. (2020). Restriction of HIV-1 escape by a highly broad and potent neutralizing antibody. Cell 180, 471–489.e22.

Seydoux, E., Homad, L.J., MacCamy, A.J., Parks, K.R., Hurlburt, N.K., Jennewein, M.F., Akins, N.R., Stuart, A.B., Wan, Y.-H., Feng, J., et al. (2020). Analysis of a SARS-CoV-2-infected individual reveals development of potent neutralizing antibodies with limited somatic mutation. Immunity 53, 98–105.e5.

Shang, J., Ye, G., Shi, K., Wan, Y., Luo, C., Aihara, H., Geng, Q., Auerbach, A., and Li, F. (2020). Structural basis of receptor recognition by SARS-CoV-2. Nature 581, 221–224.

Smith, D.J., Lapedes, A.S., de Jong, J.C., Bestebroer, T.M., Rimmelzwaan, G.F., Osterhaus, A.D.M.E., and Fouchier, R.A.M. (2004). Mapping the antigenic and genetic evolution of influenza virus. Science 305, 371–376.

Starr, T.N., Greaney, A.J., Hilton, S.K., Ellis, D., Crawford, K.H.D., Dingens, A.S., Navarro, M.J., Bowen, J.E., Tortorici, M.A., Walls, A.C., et al. (2020). Deep mutational scanning of SARS-CoV-2 receptor binding domain reveals constraints on folding and ACE2 binding. Cell 182, 1295–1310.e20.

Steffen, T.L., Taylor Stone, E., Hassert, M., Geerling, E., Grimberg, B.T., Espino, A.M., Pantoja, P., Climent, C., Hoft, D.F., George, S.L., et al. (2020). The receptor binding domain of SARS-CoV-2 spike is the key target of neutralizing antibody in human polyclonal sera.

Tian, X., Li, C., Huang, A., Xia, S., Lu, S., Shi, Z., Lu, L., Jiang, S., Yang, Z., Wu, Y., et al. (2020). Potent binding of 2019 novel coronavirus spike protein by a SARS coronavirus-specific human monoclonal antibody. Emerg. Microbes Infect. 9, 382–385.

Tong, S., Conrardy, C., Ruone, S., Kuzmin, I.V., Guo, X., Tao, Y., Niezgoda, M., Haynes, L., Agwanda, B., Breiman, R.F., et al. (2009). Detection of novel SARS-like and other coronaviruses in bats from Kenya. Emerg. Infect. Dis. 15, 482–485.

Walls, A.C., Fiala, B., Schäfer, A., Wrenn, S., Pham, M.N., Murphy, M., Tse, L.V., Shehata, L., O’Connor, M.A., Chen, C., et al. (2020a). Elicitation of potent neutralizing antibody responses by designed protein nanoparticle vaccines for SARS-CoV-2. bioRxiv 2020.08.11.247395.

Walls, A.C., Park, Y.-J., Tortorici, M.A., Wall, A., McGuire, A.T., and Veesler, D. (2020b). Structure, function, and antigenicity of the SARS-CoV-2 spike glycoprotein. Cell 181, 281–292.e6.

Wec, A.Z., Bornholdt, Z.A., He, S., Herbert, A.S., Goodwin, E., Wirchnianski, A.S., Gunn, B.M., Zhang, Z., Zhu, W., Liu, G., et al. (2019). Development of a human antibody cocktail that deploys multiple functions to confer pan-ebolavirus protection. Cell Host Microbe 25, 39–48.e5.

Wec, A.Z., Wrapp, D., Herbert, A.S., Maurer, D.P., Haslwanter, D., Sakharkar, M., Jangra, R.K., Dieterle, M.E., Lilov, A., Huang, D., et al. (2020). Broad neutralization of SARS-related viruses by human monoclonal antibodies. Science 369, 731–736.

Weisblum, Y., Schmidt, F., Zhang, F., DaSilva, J., Poston, D., Lorenzi, J.C.C., Muecksch, F., Rutkowska, M., Hoffmann, H.-H., Michailidis, E., et al. (2020). Escape from neutralizing antibodies by SARS-CoV-2 spike protein variants. bioRxiv 2020.07.21.214759.

Weissman, D., Alameh, M.-G., LaBranche, C.C., Edwards, R.J., Sutherland, L., Santra, S., Mansouri, K., Gobeil, S., McDanal, C., Pardi, N., et al. (2020). D614G Spike mutation increases SARS CoV-2 susceptibility to neutralization (medRxiv).

Wong, A.H.M., Tomlinson, A.C.A., Zhou, D., Satkunarajah, M., Chen, K., Sharon, C., Desforges, M., Talbot, P.J., and Rini, J.M. (2017). Receptor-binding loops in alphacoronavirus adaptation and evolution. Nat. Commun. 8, 1735.

Wrapp, D., Wang, N., Corbett, K.S., Goldsmith, J.A., Hsieh, C.-L., Abiona, O., Graham, B.S., and McLellan, J.S. (2020). Cryo-EM structure of the 2019-nCoV spike in the prefusion conformation. Science 367, 1260–1263.

Wu, Y., Wang, F., Shen, C., Peng, W., Li, D., Zhao, C., Li, Z., Li, S., Bi, Y., Yang, Y., et al. (2020). A noncompeting pair of human neutralizing antibodies block COVID-19 virus binding to its receptor ACE2. Science 368, 1274–1278.

Yang, X.-L., Hu, B., Wang, B., Wang, M.-N., Zhang, Q., Zhang, W., Wu, L.-J., Ge, X.-Y., Zhang, Y.-Z., Daszak, P., et al. (2016). Isolation and characterization of a novel bat coronavirus closely related to the direct progenitor of Severe Acute Respiratory Syndrome Coronavirus. J. Virol. 90, 3253–3256.

Yuan, M., Wu, N.C., Zhu, X., Lee, C.-C.D., So, R.T.Y., Lv, H., Mok, C.K.P., and Wilson, I.A. (2020). A highly conserved cryptic epitope in the receptor binding domains of SARS-CoV-2 and SARS-CoV. Science 368, 630–633.

Zhou, H., Chen, X., Hu, T., Li, J., Song, H., Liu, Y., Wang, P., Liu, D., Yang, J., Holmes, E.C., et al. (2020a). A novel bat coronavirus closely related to SARS-CoV-2 contains natural insertions at the S1/S2 cleavage site of the Spike protein. Curr. Biol. 30, 2196–2203.e3.

Zhou, P., Yang, X.-L., Wang, X.-G., Hu, B., Zhang, L., Zhang, W., Si, H.-R., Zhu, Y., Li, B., Huang, C.-L., et al. (2020b). A pneumonia outbreak associated with a new coronavirus of probable bat origin. Nature 579, 270–273.

Zhou, T., Tsybovsky, Y., Olia, A.S., Gorman, J., Rapp, M.A., Cerutti, G., Katsamba, P.S., Nazzari, A., Schon, A., Wang, P.D., et al. (2020c). A pH-dependent switch mediates conformational masking of SARS-CoV-2 spike. bioRxiv 2020.07.04.187989.

Zost, S.J., Gilchuk, P., Case, J.B., Binshtein, E., Chen, R.E., Nkolola, J.P., Schäfer, A., Reidy, J.X., Trivette, A., Nargi, R.S., et al. (2020a). Potently neutralizing and protective human antibodies against SARS-CoV-2. Nature 584, 443–449.

Zost, S.J., Gilchuk, P., Chen, R.E., Case, J.B., Reidy, J.X., Trivette, A., Nargi, R.S., Sutton, R.E., Suryadevara, N., Chen, E.C., et al. (2020b). Rapid isolation and profiling of a diverse panel of human monoclonal antibodies targeting the SARS-CoV-2 spike protein. Nat. Med.

